# Distinct allosteric mechanisms of first-generation MsbA inhibitors

**DOI:** 10.1101/2021.05.25.445681

**Authors:** François A. Thélot, Wenyi Zhang, KangKang Song, Chen Xu, Jing Huang, Maofu Liao

**Affiliations:** Department of Cell Biology, Blavatnik Institute, Harvard Medical School, Boston MA, USA; Biological and Biomedical Sciences Program, Harvard University, Cambridge MA, USA; Key Laboratory of Structural Biology of Zhejiang Province, Westlake University, Hangzhou, China; Westlake AI Therapeutics Lab, Westlake Laboratory of Life Sciences and Biomedicine, Hangzhou, China; Department of Biochemistry and Molecular Pharmacology, University of Massachusetts Medical School, Worcester MA, USA; Cryo-EM Core Facility, University of Massachusetts Medical School, Worcester MA, USA

## Abstract

Present in all kingdoms of life, ATP-binding cassette (ABC) transporters couple ATP hydrolysis to mechanical force and facilitate trafficking of diverse substrates across biological membranes. Although many ABC transporters are promising drug targets, their mechanisms of regulation by small molecule inhibitors remain largely unknown. Herein, we used the lipopolysaccharide (LPS) flippase MsbA, a prototypical ABC exporter, as a model system to probe mechanisms of allosteric modulation by compounds binding to the transmembrane domains (TMDs). Recent chemical screens have identified intriguing LPS transport inhibitors targeting MsbA: the ATPase stimulator TBT1 and the ATPase inhibitor G247. Despite preliminary biochemical and structural data, it is unclear how TBT1 and G247 bind to the MsbA TMDs yet induce opposite allosteric effect in the nucleotide-binding domains (NBDs). Through single-particle EM, mutagenesis and activity assay, we show that TBT1 and G247 bind adjacent yet separate locations in the TMDs, inducing drastic changes in TMD conformation and NBD positioning. Two TBT1 molecules asymmetrically occupy the LPS binding site to break the symmetry of MsbA, resulting in disordered transmembrane helices and decreased NBD distance. In this novel inhibited ABC transporter state, decreased distance between the NBDs causes stimulation of ATP hydrolysis yet LPS transport blockage. In contrast, G247 acts as a TMDs wedge, symmetrically increasing NBD separation and preventing conformational transition of MsbA. Our study uncovers the distinct mechanisms of the first-generation MsbA-specific inhibitors and demonstrates that rational design of substrate-mimicking compounds can be exploited to develop useful ABC transporter modulators.

## Introduction

ATP-binding cassette (ABC) transporters are a large family of integral membrane proteins capable of coupling ATP hydrolysis to substrate transport. Throughout the tree of life, ABC transporters perform diverse functions including translocating lipids across cell membranes^1–3^, mediating ion channel opening^4,5^, and exporting cytotoxic compounds^6–8^. All ABC transporters have two transmembrane domains (TMDs), which directly interact with transported substrates, and two nucleotide-binding domains (NBDs), which bind and hydrolyze ATP to provide energy for substrate translocation. Decades of biochemical and structural investigation have led to the emergence of the alternating access model^9–13^, which describes how ABC transporters couple substrate translocation to ATP hydrolysis: 1) the substrate is first recognized by the inward-facing transporter; 2) binding of ATP molecules in the NBDs promotes a transition to the outward facing state and release of the substrate across the membrane; 3) ATP hydrolysis then initiates a reset towards the inward-facing state. Interrupting such conformational transition cycle has direct applications in treating cancer^14^, regulating cholesterol homeostasis^15^, and developing new antibiotics^16^.

Many small molecule inhibitors that interfere with ABC transporter function were developed against human multidrug transporters, including the ABCB1 inhibitors zosuquidar^8,17,18^, tariquidar^8^ and elacridar^8^, as well as the ABCG2 inhibitors MZ29^19^ and MB136^19^. These compounds bind to the TMDs, interrupting conformational transition from inward to outward-facing and consequently decreasing ATPase activity^8,19,20^. Despite the wealth of inhibitor-bound structures of ABCB1 and ABCG2, very little structural data is available for most other ABC transporters, which demonstrate much narrower substrate specificity. Several critical questions remain unaddressed. Are small-molecule inhibition mechanisms transporter-specific, or shared across the ABC superfamily? Are there generally druggable conformations, domains or pockets in ABC transporters? Can ATP hydrolysis be decoupled from substrate transport as a mode of inhibition?

MsbA has long been used as a model ABC transporter^21,22^, mainly because it is biochemically well-behaved and plays an essential role in lipopolysaccharide (LPS) biogenesis in Gram-negative bacteria. Inhibition of MsbA leads to accumulation of LPS intermediates in the inner membrane, which is toxic and leads to cell death^23^, highlighting the potential of MsbA as a target for development of novel antibiotics against multidrug-resistant pathogens. High-resolution views of the conformational cycle of MsbA have been well documented by intense structural investigations using both X-ray crystallography^22,24,25^ and cryo-EM^1,26^. Much progress has been achieved: the full protein structure is now known in several conformations including ligand-free^22,24^, LPS-bound^1,25^, adenylyl-imidodiphosphate (AMP-PNP)-bound^22^ and vanadate-trapped^1,22^. Importantly, the basis of substrate specificity for MsbA has been revealed — the LPS binding pocket is formed by a ring of basic residues and a large hydrophobic pocket, which interact with the phosphorylated glucosamines and acyl chains in LPS, respectively^1,25,27^.

In contrast to the rapid progress in understanding the detailed mechanism of MsbA-driven LPS flipping, investigation on small molecule inhibition of MsbA has lagged behind, hindering the discovery of antibiotics to block LPS transport and outer membrane biogenesis^27^. To date, only two kinds of MsbA-specific inhibitors have been reported, demonstrating distinct effects on MsbA. One inhibitor with tetrahydrobenzothiophene (TBT) scaffold, named herein as TBT1, stimulates MsbA ATPase activity despite abolishing LPS transport^28^. Such decoupling of ATPase activity from substrate transport is extraordinary, because the vast majority of ABC transporter inhibitors have a repressive effect on ATP hydrolysis^12^. Analysis of TBT1-resistant mutants suggests that the compound binds in the TMDs of MsbA, though the mechanism of action of TBT1 is unclear. The other class of MsbA inhibitors, here referred to as G compounds, was identified in a high-throughput chemical screen and shown to block both ATP hydrolysis and substrate transport^25,29^. Crystal structures of MsbA in complex with two compounds of this class, G907 and G092, reveal an inhibitor binding pocket in the upper TMDs and a conformational change characterized by a narrowed asymmetric positioning of the NBDs. Despite previous pioneering structural studies^25^, it remains puzzling how inhibitor binding in two symmetry-related, identical pockets induces asymmetric conformational change in the homodimeric MsbA. In summary, while these first-generation inhibitors pave the way to antibiotic development targeting MsbA-dependent LPS transport, how these small molecules block LPS transport while exerting opposite allosteric effects on ATP hydrolysis is largely elusive.

In this work, we have exploited single-particle cryo-EM to study how TBT1 and a G compound (G247) differentially modulate MsbA function in a lipid bilayer environment. These compounds bind nearby pockets in the TMDs, displacing surrounding transmembrane helices (TMs) and allosterically inducing distinct NBDs positioning. TBT1 binding results in an unprecedented collapsed conformation, characterized by highly disordered domain-swapping TMs and drastically reduced inter-NBD spacing. This is a unique case in which two copies of a small molecule mimic the much larger LPS substrate and induce asymmetrical conformational changes in MsbA. Virtual screening against the TBT1 pocket resulted in the identification of a novel scaffold with similar ATPase stimulatory effect, demonstrating the feasibility of targeting ABC transporter induced-fit substrate binding pockets for modulator development. Meanwhile, our analyses formally proved that G247 functions as a TMD wedge that symmetrically widens NBD spacing and prevents transporter closure and transition to the outward-facing state. Taken together, our results show that MsbA modulators induce distinctive conformations and suggest inter-NBD spacing as a heuristic for classifying ATPase stimulators from inhibitors.

### TBT1-bound MsbA adopts an asymmetric, collapsed inward-facing conformation

TBT1 was identified as an LPS transport inhibitor and MsbA ATPase stimulator in strains from the *Acinetobacter* genus^28^. We therefore sought to explore the mechanism of TBT1 inhibition in *Acinetobacter baumannii*, an ESKAPE^30^ pathogen responsible for antibiotic-resistant infections in patients. *A. baumannii* MsbA was expressed in *Escherichia coli* cells, purified in dodecyl maltoside (DDM) and reconstituted in palmitoyl-oleoyl-phosphatidylglycerol (POPG) nanodiscs^31,32^ (Suppl. Fig. 1a-f). Basal ATPase activity of *A. baumannii* MsbA in nanodiscs was ~1 μmol ATP/min/mg MsbA (Fig. 1a), which is ~4-fold lower than *E. coli* MsbA^1^ but considerably more active than when previously characterized in liposomes^28^. Exposure of MsbA to TBT1 resulted in ~4-fold stimulation in ATPase activity (Suppl. Fig. 1d), as well as dose-dependent increase in reaction rate in Michaelis-Menten kinetics (Fig. 1a). After establishing lipid nanodisc as a suitable system for MsbA stimulation by TBT1, we used single-particle cryo-EM to determine the structure of TBT1 -bound MsbA at an overall resolution of 4.3 Å, with the TMDs at 4.0-Å resolution showing sufficient side-chain densities for model building (Fig. 1b and Suppl. Fig. 2,3).

**Fig. 1.**
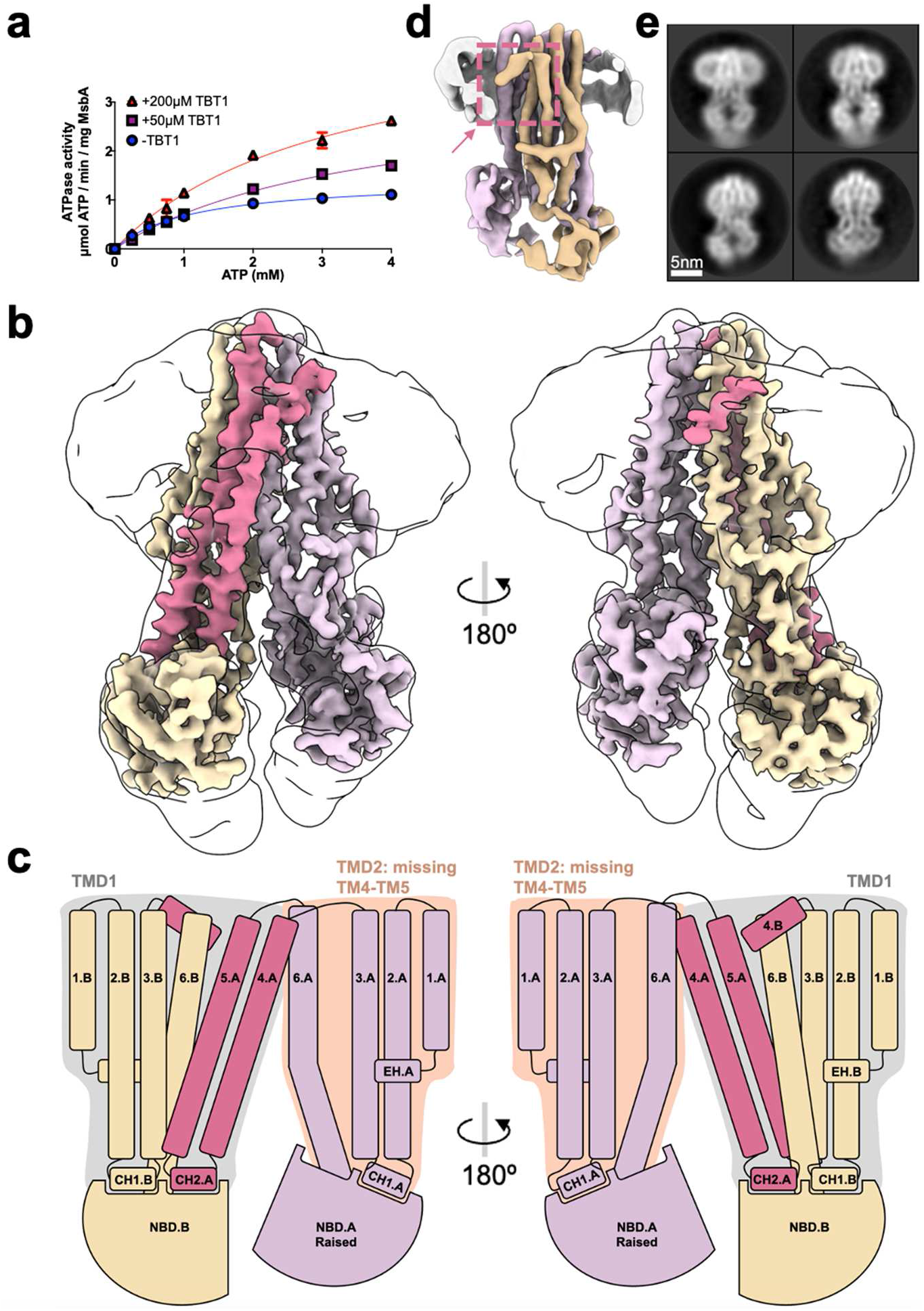
TBT1 binding induces a collapsed inward-facing conformation of MsbA. **a,** ATPase activity of *A. baumannii* MsbA in nanodiscs, measured at varying ATP and TBT1 concentrations. TBT1 increases the rate of ATP hydrolysis in a dose-dependent manner. Each point represents mean ±SD (n=3). **b,** 4.0-Å resolution cryo-EM map of *A. baumannii* MsbA in complex with TBT1, with domain-swapping TM4-TM5 colored fuchsia. The unsharpened map filtered at 10-Å resolution is displayed as outline to show the nanodisc and tightened NBDs positioning. **c,** Cartoon of TBT1-bound MsbA with structural elements indicated, illustrating the structural asymmetry of the two MsbA chains. EH, elbow helix; CH, coupling helix. **d,** Cryo-EM reconstruction low-pass filtered at 6-Å resolution. The dashed box points to the N-terminal end of TM4.B, where TM4-TM5.B becomes disordered and absent from the reconstruction. **e,** Representative 2D class averages of TBT1-bound MsbA, showing constricted TMDs yet separate NBDs. Box size is 203 Å.

The conformation of TBT1-bound MsbA is asymmetric and distinct from all previously determined structures of type IV ABC transporters^33^ that are characterized by their domain-swapped TMs. Notably, the domain-swapping TM4-TM5 bundle is well resolved in chain A yet disordered in chain B (Fig. 1b,c), which is evident also in unmasked and unsharpened 3D reconstructions (Suppl. Fig. 4b). This chain mismatch results in asymmetrical TMDs, with TMD1 including 6 helices (TM4,5.A, TM1,2,3,6.B) and TMD2 including only 4 helices (TM1,2,3,6.A). While TM4.B is mostly unresolved, its N-terminal end is clearly discernable and forms a 61° angle with the C-terminal end of TM3.B, which is ~15° greater than the corresponding angle between the well resolved TM4.A and TM3.A (Suppl. Fig. 4c). Accordingly, TM4.B kinks outwards, pushing the nanodisc density to bulge upwards from the membrane plane (Fig. 1d and Suppl. Fig. 4c, left). The domain-swapping TM4-TM5 bundle is of critical importance because the coupling helix (CH) at the TM4-TM5 junction (CH2), together with CH1 at the TM2-TM3 junction, is responsible for transmitting movement from the catalytically active NBDs to the TMDs (Fig. 1c). Due to disordered TM4-TM5.B, CH2.B is absent for interaction with the NBD in chain A (NBD.A). Accordingly, NBD.A tilts up, moving towards the NBD dimerization interface, and becomes more mobile as its density is considerably weakened compared to that of the opposing NBD.B (Fig. 1b,c). Notably, CH1.A also appears destabilized, likely because it cannot form proper interaction with the drastically shifted NBD.A (Suppl. Fig. 4d). In summary, the raised NBD.A is disengaged from its two coupling helices and seems decoupled from the TMDs (Suppl. Fig. 4d).

At first sight, 2D class averages and 3D reconstruction of the cryo-EM images of TBT1-bound MsbA (Fig. 1e and Suppl. Fig. 4e) resemble those of nucleotide-bound MsbA^1^, both exhibiting constricted TMDs and reduced inter-NBD distance compared to nucleotide-free MsbA. However, the TBT1-bound MsbA conformation is still inward-facing because no nucleotide is present in the cryo-EM sample and the NBDs remain fully separate. Accordingly, TMD1 is well aligned to the structure of nucleotide-free *E. coli* MsbA (Suppl. Fig. 4f, root-mean-square-deviation (RMSD) of 5.5 Å over 314 Cα atoms), leaving significant structural mismatch from TMD2. The most noteworthy structural rearrangement in TMD2 is the 13-Å displacement of TM6.A towards the central cavity (Suppl. Fig. 4g), which seems to pull NBD.A upwards and to close proximity of NBD.B. Thus, the TBT1-bound MsbA structure described herein represents an unusual “collapsed inward-facing” conformation, which is characterized by two unprecedented features: 1) complete destabilization of one domain-swapping TM4-TM5 helix bundle; 2) highly asymmetric positioning of the NBDs.

### TBT1 binding drives the closure of a wide open MsbA

To determine the exact conformational changes of MsbA upon TBT1 binding, we sought to characterize the structure of drug-free *A. baumannii* MsbA. To this end, we imaged *A. baumannii* MsbA in identical conditions as described above, but without addition of TBT1. The resulting cryo-EM reconstruction at an overall resolution of 5.2 Å enabled unambiguous tracing of each TM helix, revealing a wide inward-facing conformation (Fig. 2a, Suppl. Fig. 5). To our knowledge, this is the first structure of membrane embedded MsbA in a wide inward-facing conformation. Such conformation had only been characterized by X-ray crystallography using MsbA in detergents^21,24^, and its physiological relevance has been heavily debated^1,26^. Our result demonstrates that the wide inward-facing conformation of MsbA can exist in a lipid bilayer and may not be an artifact caused by deprivation of membrane environment and the usage of detergents. Furthermore, the wide opening of MsbA seems not due to lack of bound substrate, because a clear LPS-like density is present in the inner cavity (Suppl. Fig. 5g). This is consistent with previous reports that *A. baumannii* MsbA, which transports endogenous LPS with 7 acyl chains, is competent to bind and flip *E. coli* LPS containing 6 acyl chains^28^.

**Fig. 2.**
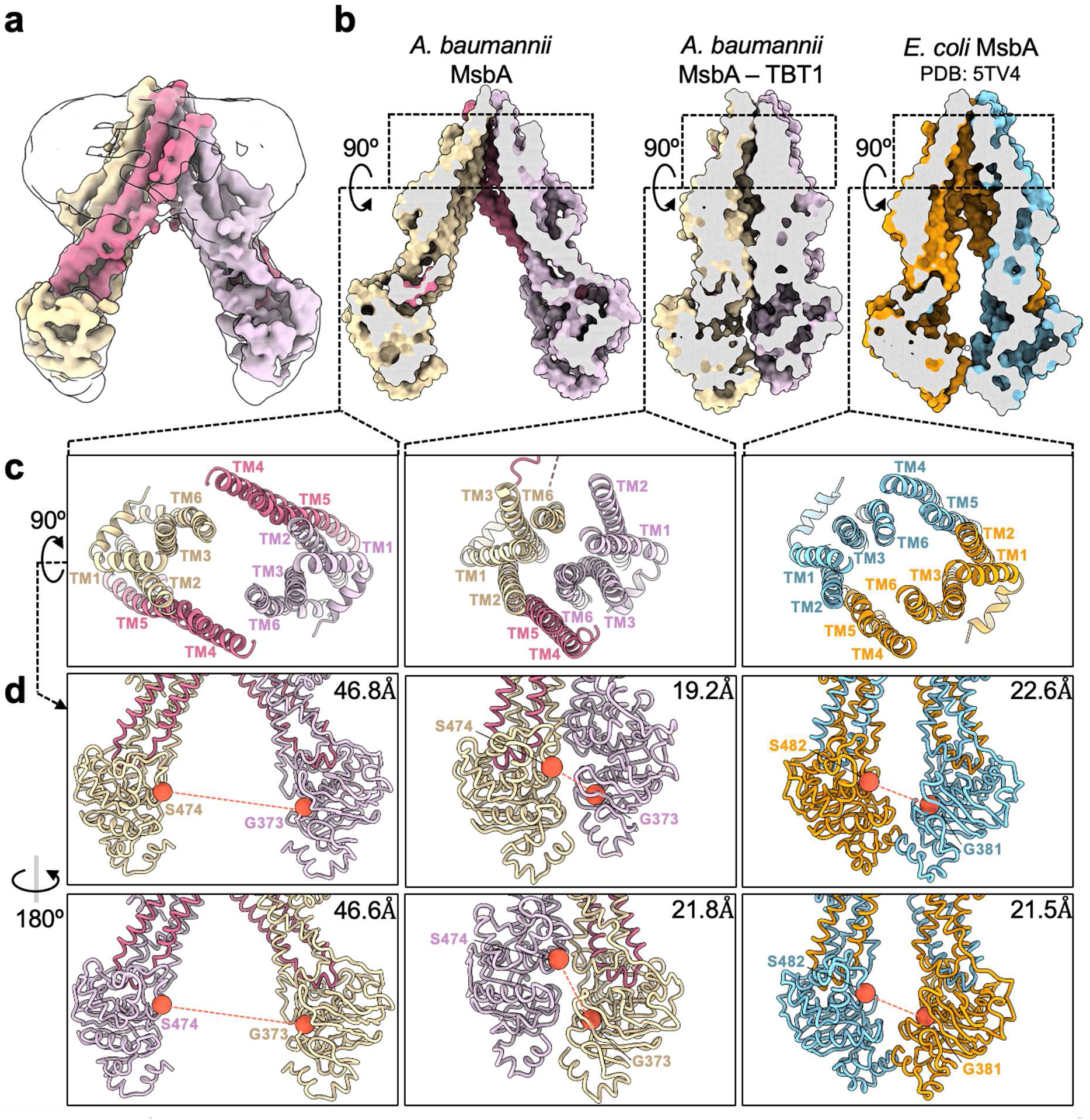
Drug-free *A. baumannii* MsbA in nanodiscs adopts a wide inward-facing conformation. **a,** 5-Å resolution cryo-EM reconstruction of *A. baumannii* MsbA in the absence of ligand. The unsharpened map filtered at 8-Å resolution is displayed as outline to show the nanodisc. **b,** Section through surface representation and comparison for three structures: A. *baumannii* MsbA, TBT1-bound A. *baumannii* MsbA, and *E. coli* MsbA (PDB: 5TV4). **c,** Top-down view of the substrate binding pockets. **d,** Two side views of the NBDs. Cα distances between ABC signature motif serine and opposing Walker A glycine residues are indicated as dashed lines, with values shown in each panel.

Since drug-free *A. baumannii* MsbA has well defined domain-swapping TM4-TM5 bundle in both TMDs and distanced NBDs, TBT1 binding is solely responsible for converting the wide open MsbA to the collapsed inward-facing conformation. To gain insights of TBT1 stimulated ATP turnover of MsbA, we compared three cryo-EM structures: TBT1-bound or drug-free *A. baumannii* MsbA, and *E. coli* MsbA (Fig. 2b-d). Notably, the central substrate-binding pocket of TBT1-bound MsbA is much more constricted than that in the other two structures (Fig. 2b,c), consistent with the notion that the TMDs in TBT1-bound state present similarities with an outward-facing transporter. Upon TBT1 binding, the NBD distance is drastically decreased from ~47 Å to ~20 Å and even slightly shorter than the NBD distance in LPS-bound *E. coli* MsbA (Fig. 2d). The two NBDs of TBT1-bound MsbA are positioned asymmetrically, such that one ATP binding site is tightened compared to the opposite ATP site (19.2 Å vs. 21.8 Å, Fig. 2d). The structural analysis of TBT1-bound MsbA provides an intuitive explanation for ATPase activity stimulation: removal of TM4-TM5.B bundle and sliding of TM6.A into the central cavity greatly reduce inter-NBD distance, thus increasing the speed of NBD dimerization and ATP hydrolysis.

### TBT1 hijacks the LPS binding site to modulate MsbA

Two densities, each consistent with a TBT1 molecule, are present in the upper region of the TMDs (Fig. 3a,b, Suppl. Fig. 6a,b). Interestingly, these two TBT1 ligands are positioned asymmetrically in the central substrate pocket of MsbA and related by a ~60° rotation angle, with the carboxyl group pointing downwards to the cytoplasm. The general binding mode of TBT1 is reminiscent of how *E. coli* MsbA binds the amphiphilic LPS using a large hydrophobic pocket, which accommodates the lipid acyl chains in LPS, and a ring of basic residues (Arg78, Arg148 and Lys299), which stabilize the phosphate groups on glucosamines^1^. Similarly, the two TBT1 molecules are located right above the ring of basic residues, with the aromatic ring structures of TBT1 positioned in the hydrophobic pocket (Fig. 3d).

**Fig. 3.**
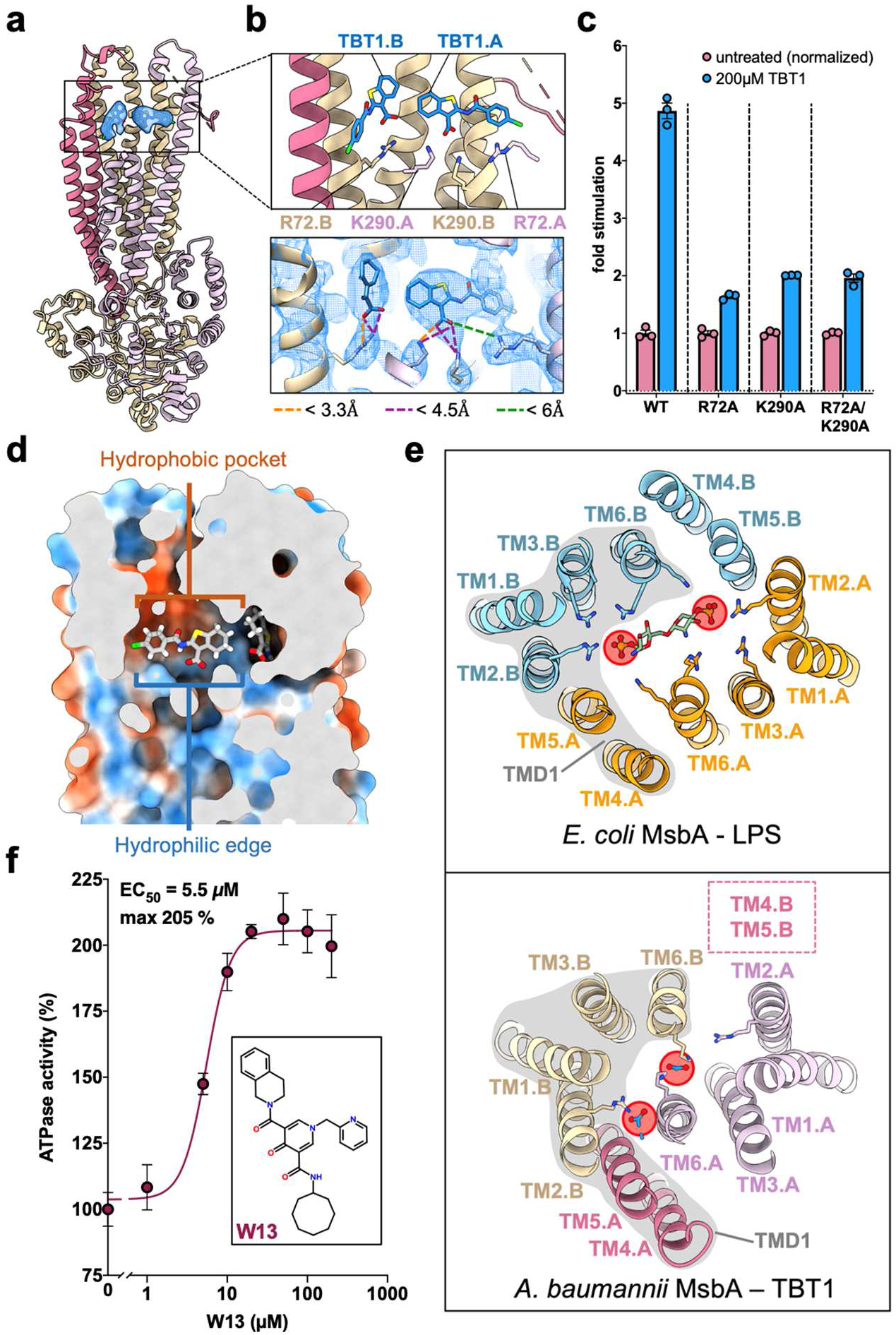
TBT1 binding to MsbA. **a,** Cartoon representation of TBT1-bound *A. baumannii* MsbA colored as in Fig. 1b, with TBT1 colored blue. **b,** Close-up view of the TBT1 binding site (top) and superimposition of model and cryo-EM density (bottom). Distances between TBT1 carboxyl group and neighboring lysine and arginine residues are shown as colored dashed lines. **c,** TBT1-induced ATPase stimulation of wild-type and mutant MsbA. The activity of each protein was normalized to its basal activity without TBT1. Error bars correspond to mean ±SD (n=3). **d,** Surface representation of the TBT1 binding pocket, with hydrophilic and hydrophobic surfaces colored blue and orange, respectively. **e,** Top-down view of *E. coli* MsbA bound to LPS (PDB: 5TV4) and *A. baumannii* MsbA bound to TBT1. TMD1 is conformationally similar in both structures and indicated with a gray background. The phosphate groups on LPS glucosamines and the carboxyl groups of TBT1 are marked with red circles. **f,** ATPase activity of *A. baumannii* MsbA with W13 at increasing concentrations. Each point represents mean ±SD (n=3). The molecular structure of W13 is shown in inset.

Despite being located in asymmetric binding pockets (Suppl. Fig. 6b), each TBT1 molecule is stabilized by an analogous set of hydrogen-bonding and electrostatic interactions between TBT1 carboxyl groups and neighboring basic sidechains. TBT1.A is oriented parallel to the membrane plane, with its carboxyl group within hydrogen-bonding distance of Lys290.A and forming long-range electrostatic interactions with Lys290.B and Arg72.A, and TBT1.B is rotated relative to TBT1.A, forming a salt bridge with Arg72.B (Fig. 3b). Strikingly, the TBT1 densities and neighboring residues have better-defined features than the rest of the TMDs, suggesting that TBT1 binding stabilizes the surrounding pocket (Fig. 3b, Suppl. Fig. 6a).

Our results suggest that the carboxyl group of TBT1 is analogous to the phosphate groups on LPS glucosamines, both forming electrostatic interaction with MsbA. We thus sought to mutate Arg72 and Lys290 to determine whether MsbA retains sensitivity to TBT1-induced ATPase stimulation. *A. baumannii* MsbA with R72A, K290A, or R72A/K290A mutation was purified as wild-type MsbA and reconstituted in nanodiscs (Suppl. Fig. 6c,d). All MsbA mutants demonstrated largely diminished TBT1-induced stimulation compared to the wild-type protein, highlighting the importance of electrostatic interactions in TBT1 binding. Consistently, chemical modification of the carboxyl group of TBT1 renders the compound incompetent for MsbA inhibition^28^. Although the electrostatic interactions described herein may be the main drivers of TBT1 binding, hydrophobic contacts through the ring structures of TBT1 also participate in stabilizing the inhibitor. For instance, L68F and L150V mutants, which were previously shown to confer resistance to TBT1^28^, are located in the vicinity of TBT1 binding sites (Suppl. Fig. 6e). While not directly adjacent to TBT1, it is conceivable that these mutations allosterically alter the pocket and prevent efficient inhibitor binding.

Despite striking resemblance in LPS and TBT1 recognition by MsbA, differences in TM positioning lead to divergence in ligand binding (Fig. 3e). LPS and TBT1 stabilization both involve TM2 (Arg72) and TM6 (Lys290), although unlike LPS, TBT1 does not clearly interact with TM3. Notably, the two TBT1 copies are a distorted mimic of LPS, with decreased distance between TBT1 carboxyl groups compared to LPS glucosamine phosphates (Fig. 3e). Upon TBT1 binding, TMD2 (TM1,2,3,6.A) undergoes significant changes: TM6.A moves into the center of the inner cavity, presenting Lys290.A for interaction with the hydroxyl group of TBT1.A and dragging the connected NBD.A towards the central axis (Fig. 3e, bottom). TMD2 then collapses around TM6.A, with TM2.A moving closer to TM6.B. It is conceivable that the convergence between TM2.A and TM6.B would then push TM4-TM5.B laterally. The unsolvable contradiction between NBD.A moving towards the central axis and TM4-TM5.B being pinched outwards results in the dissociation of the CH2.B coupling helix from NBD.A and to the peculiar disordered domain-swapped helices of the collapsed inward-facing MsbA conformone NBD towards the dimerization interfaceation.

To our knowledge, inhibition by TBT1 represents the first example of a small molecule inhibitor targeting the central binding site of a non-multidrug ABC transporter. Because most ABC transporters have relatively narrow substrate spectra, our findings suggest that mimicking substrate binding can be a generally applicable strategy for developing small molecule modulators for all ABC transporters. To test if the central pocket is useful for structure-based compound discovery, we performed an *in silico* chemical library screen against the TBT1 induced-fit pocket. Virtual screening of ~800,000 compounds resulted in the identification of 17 hits. One of these molecules, W13, stimulated ATPase activity with an EC50 lower than TBT1 (~5.5 μM vs 13 μM, Fig. 3f and Suppl. Fig. 7b). W13 is structurally dissimilar to TBT1 and considerably larger in size (499 Da vs. 336 Da), and a single W13 molecule is expected to occupy the central MsbA pocket (Suppl. Fig. 7c,d). Molecular dynamics simulation of W13-bound MsbA in a POPC lipid bilayer suggests a reminiscent binding mode to TBT1, including hydrophobic groups of W13 extending upward in the hydrophobic pocket and carbonyl groups of W13 in close proximity to TBT1-recognizing arginine and lysine residues (Suppl. Fig. 7d).

### G247 symmetrically increases inter-NBD spacing and prevents MsbA closure

Distinct from TBT1, G compounds are the only other class of known MsbA-specific inhibitors and were shown to bind the inward-facing MsbA conformation and prevent ATP hydrolysis^25^. Crystal structures of G compound-bound *E. coli* MsbA in FA-3 detergent present asymmetrical NBDs positioning^25^, although it remains unclear how the asymmetric conformation translates to impaired conformational cycling. Furthermore, the extraordinary conformational flexibility of ABC transporters can result in different states being captured depending on experimental conditions, a phenomenon which has been observed with the apo MsbA being wide-open in crystal structures^21,22,24^ yet adopting a narrow inward-facing conformation^1,26^ in cryo-EM reconstructions. This emphasizes the importance of comparing different conformations of ABC transporters under as close to identical conditions as is possible. In this context, side-by-side comparison of cryo-EM structures of nanodisc-embedded G inhibitor-bound MsbA and drug-free MsbA^1^ would provide unbiased insight into their inhibition mechanism.

We obtained the cryo-EM map of nanodisc-embedded *E. coli* MsbA in complex with one of the G compounds (G247) at an overall resolution of ~4.1 Å (3.9 Å TMD, Fig. 4a and Suppl. Fig. 8b). The G247 density is well resolved in the upper TMDs, near the TBT1 and LPS binding sites (Fig. 4b,c), and is consistent with the G compounds in previous crystal structures^25^. As in earlier work^1,25^, LPS copurified with the protein and is bound to the central pocket (Suppl. Fig. 8c).

**Fig. 4.**
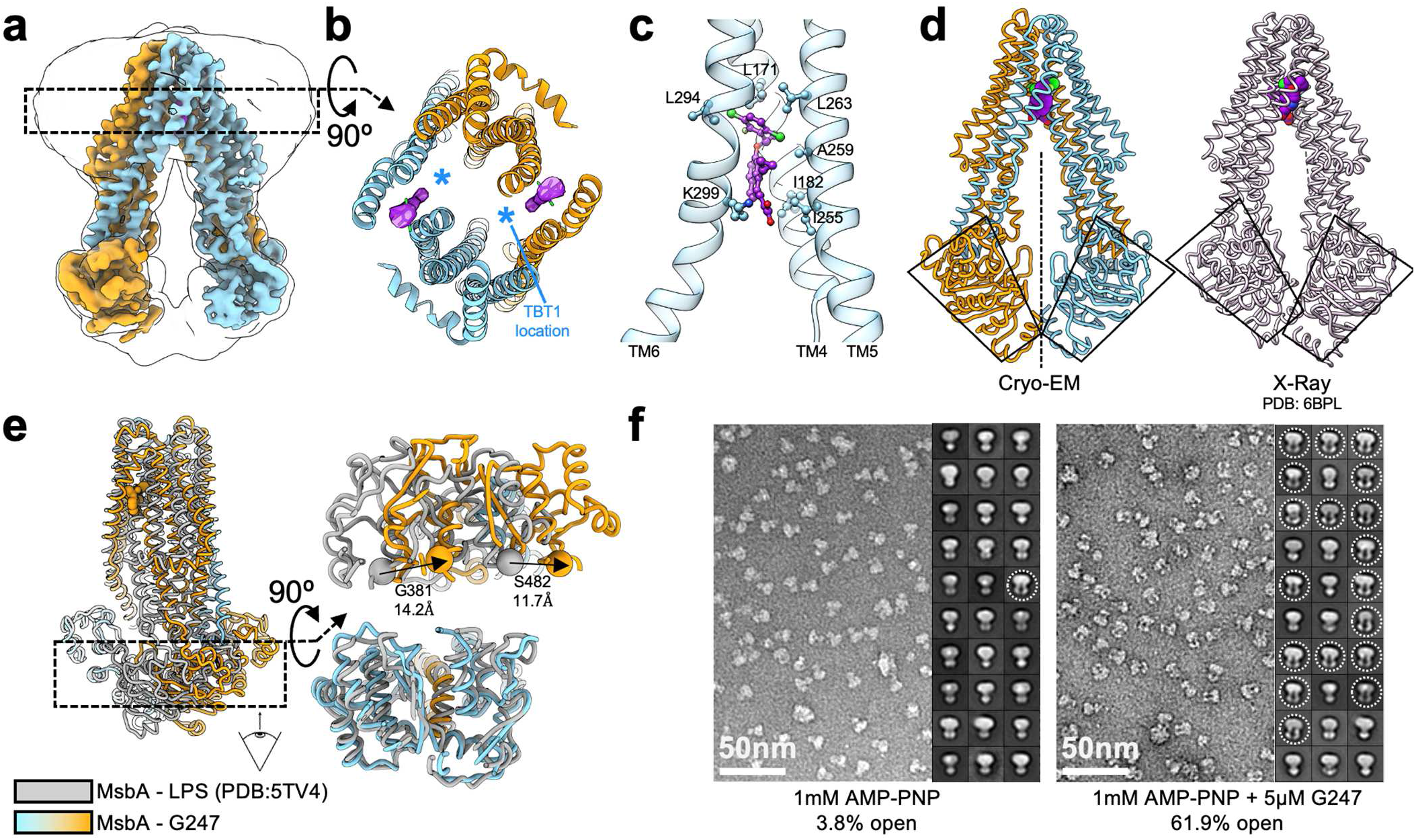
G247 symmetrically increases inter-NBD spacing and blocks MsbA closure. **a,** 3.9-Å resolution cryo-EM reconstruction of *E. coli* MsbA in complex with G247. The unsharpened map filtered at 6-Å resolution is displayed as outline to show the nanodisc. **b,** Top-down sectional view of the upper TMD region. G247 is shown purple. **c,** Side view of G247 binding pocket, viewed from the central LPS binding pocket. **d,** Comparison of MsbA structures determined by cryo-EM or X-ray crystallography (PDB: 6BPL). The cryo-EM structure adopts near-perfect C2 symmetry, while NBDs are asymmetrically positioned in the crystal structure. **e,** Comparison of the cryo-EM structures of drug-free (PDB: 5TV4, gray) and G247-bound (colored) MsbA, both in nanodiscs. G247 pushes NBDs away from each other, compared to the uninhibited state. The NBD shift is apparent when viewing MsbA from the side (left panel), and from the cytoplasmic space (right panel). **f,** Representative negative-stain EM images and 2D class averages of *E. coli* MsbA exposed to AMP-PNP, in the absence and presence of G247.

In previous crystal structures, the NBDs are brought together compared to apo state and positioned asymmetrically, such that one NBD is raised relative to the other (Fig. 4d, right). In contrast, our cryo-EM structure exhibits clear C2 symmetry, even when refined without symmetry constraints (Fig. 4d, left, and Suppl. Fig. 8d). 2D class averages of G247-bound MsbA also exhibit symmetrical NBD arrangement (Suppl. Fig. 8e), unlike TBT1-bound MsbA in which asymmetrical NBD positioning could be directly observed from 2D averages (Fig. 1e). In the asymmetric crystal structures, Arg190 on one chain forms a salt bridge with the acrylic acid tail of the G compounds while the opposite-chain Arg190 cannot be modelled^25^. It is conceivable that this transient electrostatic interaction between Arg190 and G compounds participates in stabilizing an asymmetric state with one raised NBD, which is readily captured in a constrained crystal lattice (Suppl. Fig. 8f). In contrast, Arg190 in our cryo-EM structure seems too distant from G247 to form a salt bridge, possibly accounting for the symmetrical NBD positioning observed in nanodisc (Suppl. Fig. 8g). In summary, we posit that the C2-symmetric state is the predominant conformation of G247-bound MsbA in a lipid bilayer, and that the previously characterized asymmetric NBD positioning is unlikely a major contributor to ATPase inhibition caused by G compounds.

To better understand the mechanism of action of G compounds, we aligned our cryo-EM structure of G247-bound MsbA with the crystal structure of MsbA in complex with G907^25^ (PDB:6BPL) and with the drug-free conformation^1^ (PDB:5TV4). While the G compound binding pocket outlined by TM4,5,6 is largely identical for G247 and G907 (Suppl. Fig. 8g), the positioning of TM4,5,6 relative to TM1,2,3 is different for both drug-bound structures, with a larger outward shift of TM4,5,6 for G907-bound MsbA (Suppl. Fig. 8h). Interestingly, this mismatch in TM4,5,6 positioning allosterically propagates to the rest of the TMDs and impacts the spacing between coupling helices CH1 and CH2. While CH1-CH2 spacing is reduced for G907 compared to the drug-free conformation, CH1-CH2 distance is unexpectedly increased in the G247-bound state (Suppl. Fig. 8i). Further comparison with the drug-free conformation reveals that inter-NBD distance is increased by ~13 Å upon G247 binding, with the NBDs symmetrically sliding away from each other parallel to the membrane plane (Fig. 4e). Notably, this finding is opposite to prior crystallography studies, in which G compound binding would push one NBD towards the dimerization interface by 10-15 Å^25^. In summary, while drug-free MsbA has aligned NBDs in a head-to-tail manner and thus primed for ATP hydrolysis, the increased inter-NBD distance in G247-bound MsbA is expected to reduce NBD dimerization efficiency and ATPase activity.

To formally test the hypothesis that G compounds inhibit MsbA by preventing NBD dimerization, we exposed MsbA to the non-hydrolyzable nucleotide AMP-PNP, in the presence or absence of G247 (Fig. 4f). AMP-PNP binds to the ATP sites and traps NBDs in a tightly dimerized conformation^22^, which can be readily observable in negative stain EM images. When subjected to 1mM AMP-PNP, most MsbA particles exhibit closed conformation with dimerized NBDs, with only 4% particles showing well separated NBDs. In contrast, in the presence of 5 μM of G247, many more MsbA proteins remain in inward-facing conformation, and the number of particles with separate NBDs was increased to ~62% (Fig. 4f). Taken together, our cryo-EM and negative-stain EM analyses demonstrate that G compounds act as TMD wedges, symmetrically increasing NBD separation and preventing proper conformational transition induced by ATP binding.

## Discussion

In this work, we have elucidated the distinct inhibition mechanisms of two first-generation MsbA inhibitors in a consistent nanodisc-based environment and shown that these compounds modulate ATPase activity of MsbA by varying NBD positioning. We propose the following models for the divergent action of TBT1 and G247 (Fig. 5).

i. Two copies of TBT1 asymmetrically bind in the central MsbA pocket and mimic the endogenous LPS substrate. As with LPS, TBT1 establishes both hydrophobic and electrostatic interactions with MsbA. Binding of TBT1 leads to drastic structural rearrangements predominantly within one half of the transporter, including the sliding of TM6.A into the central cavity, and destabilization of domain-swapping TM4-TM5.B bundle and coupling helices. These conformational changes result in one NBD moving towards the NBD dimerization interface. Reduction of inter-NBD distance promotes ATP hydrolysis and accounts for the stimulatory effect of TBT1.
ii. G247 binds in a pocket enclosed by TM4, TM5 and TM6, in proximity to the TBT1 and LPS binding sites. It acts as a TMD wedge and symmetrically pushes the NBDs away from each other. The increased inter-NBD spacing prevents NBD closure and results in a reduction in ATPase activity.

**Fig. 5.**
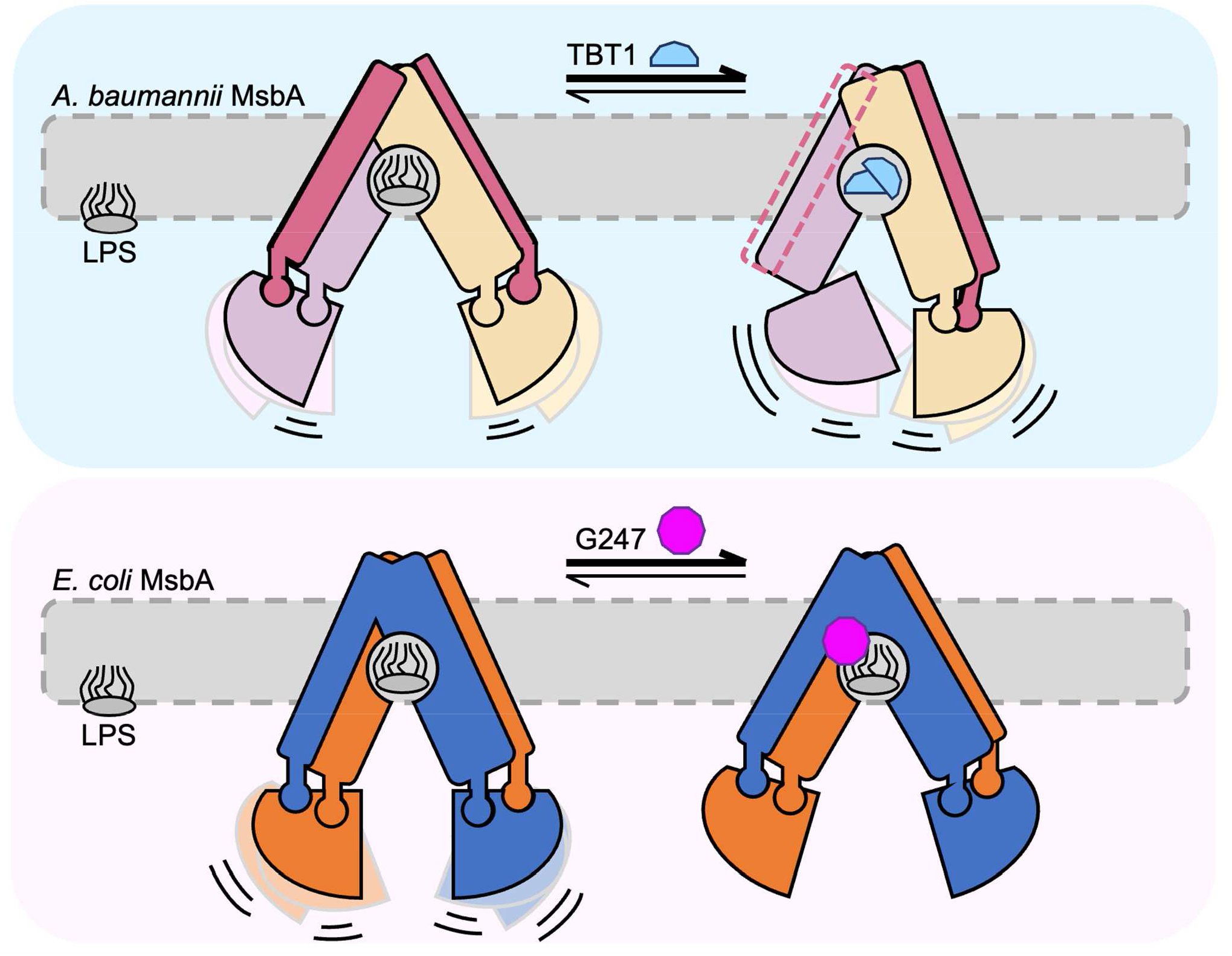
Distinct MsbA inhibition mechanisms of positive and negative allosteric ATPase modulators. See text for description of proposed mechanisms of action for TBT1 (top) and G247 (bottom). The inner membrane is shown as dashed gray box, and MsbA is colored as in previous figures. For illustrative purposes, the weakly associated CH1.A is not represented in TBT1-bound MsbA. Curved lines and duplicated NBDs illustrate NBD conformational heterogeneity and their ability to dimerize for hydrolyzing ATP, in all conformations except G247-bound MsbA.

In these inhibitor-bound structures, inter-NBD spacing compared to apo state is correlated with ATPase activity modulation. We anticipate that this simple heuristic will generalize to other inhibited ABC transporter structures.

The peculiar effect on MsbA conformation and the inhibition mechanism of TBT1 are drastically different from those of small molecule modulators for other ABC transporters, which typically induce more modest conformational changes (e.g., ABCB1 and ABCG2 in Suppl. Fig. 6f). Interestingly, the narrowed inter-NBD spacing of TBT1-bound MsbA is more akin to the NBD tightening observed when the human multidrug transporter ABCC1 is exposed to its endogenous leukotriene substrate^6^ (Suppl. Fig. 6f), further suggesting that TBT1 triggers substrate recognition-like behavior in MsbA by mimicking the much larger LPS substrate. Yet unlike the authentic ligand, TBT1 breaks the symmetry of the MsbA homodimer by destabilizing one domain-swapping helix bundle, in an unprecedented mechanism in ABC transporter family. It is noteworthy that uncoupling of ATPase function and substrate transport by TBT1 occurs through the intermediary of the domain-swapped coupling helix, which is a landmark feature of the type IV exporter fold^33^. Thus, small molecule decouplers acting like TBT1 may be identified for many other type IV exporters such as ABCB1 and ABCC1.

In contrast with TBT1, we have shown that G247 suppresses ATPase activity by increasing inter-NBD distance. This inhibition mechanism is similar to that of previously described ABC transporter inhibitors, including zosuquidar that reduces the activity of the homologous ABCB1 transporter by pushing its NBDs away from each other^17^. As the vast majority of ABC transporter inhibitors characterized so far are ATPase suppressors, it is likely that more compounds act similarly to G247 than to TBT1.

TBT1 and G247 currently face severe drawbacks preventing their application in clinical settings, including low binding affinity to MsbA^28^ and interaction with animal serum^29^. Yet compound-bound ABC transporter structures are essential tools for discovering new inhibitors, because molecules such as TBT1 and G247 reveal unexpected binding pockets by induced-fit. For example, convenient identification of W13 as an ATPase stimulator is very unlikely if virtual screening was performed against the structure of wide inward-facing, drug-free *A. baumannii* MsbA instead of TBT1-bound MsbA. We therefore hope that these ligand-bound MsbA structures will guide the development of novel classes of antibiotics targeting MsbA. One promising direction for compound optimization may consist of tethering together two TBT1-like molecules, such that inhibitor local concentration is increased upon entering the central pocket. Another worthwhile undertaking may consist in generalizing the LPS-mimicking inhibition strategy to other Gram-negative strains including antibiotic resistant pathogens. While TBT1 currently only works on *A. baumannii* MsbA, the inhibition mechanism described here should in principle extend to MsbA in other Gram-negative bacteria that presumably use a similar mechanism of LPS binding.

In summary, we have described how the first-generation MsbA-specific inhibitors TBT1 and G247 bind neighboring pockets in the TMDs yet cause opposite allosteric effects on ATPase activity through distinct mechanisms. These structures have revealed fit-induced pockets amenable for protein inhibition and should accelerate the discovery of compounds with improved affinity and deliverability.

## Acknowledgements

We are grateful to D. Kahne and D. Dates for providing the initial TBT1 sample, as well as to Genentech, Inc. for their donation of G247. We are thankful to J. Zhang for helping with ATPase screening experiments. We thank Z. Li, S. Sterling, R. Walsh and S. Rawson from Harvard cryo-EM center for Structural Biology for their training and expert advice. The majority of the datasets were collected at the Cryo-EM core facility at University of Massachusetts Medical School. We are grateful to all Liao lab members for their helpful feedback throughout this project. M.L. was supported by an NIH grant from the NIGMS (R01 GM122797).

## Author Contribution

M.L. conceived and supervised the project. F.T. performed molecular cloning, protein purification, nanodisc reconstitution, ATPase assays, negative-stain EM, sample vitrification, cryo-EM data collection and processing, and model building. K.S. and C.X. helped cryo-EM data acquisition. W.Z and J.H performed virtual screening and molecular dynamics simulation. F.T. and M.L. drafted the manuscript. All authors contributed to data analysis and manuscript preparation.

## Author Information

The authors declare no competing financial interests. Correspondence and requests for materials should be addressed to M.L.

## Materials and Methods

### Cloning, expression, and purification of MsbA

The gene encoding for *A. baumannii* MsbA was amplified from genomic DNA (ATCC strain 2208, Ref 19606) and cloned into pET-28a vector with an additional N-terminal His tag. *A. baumannii* MsbA alanine mutants were introduced by Quikchange (Agilent) mutagenesis. The *E.coli* MsbA construct used in this study was originally obtained from the G. Chang lab^1,22^ and consists of N-terminally His tagged MsbA in pET-19b. Both gene variants had similar biochemical behavior and were therefore expressed and purified in identical conditions. The MsbA constructs were both transformed in *E. coli* BL21 DE3 and grown in Terrific Broth medium to OD600 ~1.0 at 37°C, before induction with 1mM β-D-1-thiogalactopyranoside (IPTG) and overnight growth at 17°C. Cells were subsequently harvested by centrifugation, resuspended in Buffer A (50 mM Tris pH 7.8, 300 mM NaCl, 10% glycerol, 0.5 mM TCEP), and lysed with a probe sonicator. Lipid membranes were harvested by 75min ultracentrifugation at 100,000 xg at 4°C, frozen in liquid nitrogen and stored at −80°C for future use. Membranes were thawed in buffer A + 1% (w/v) n-dodecyl-β-D-maltopyranoside (DDM), resuspended with a Dounce homogenizer and stirred for ~1 hour at 4°C. Insoluble debris were harvested by centrifugation at ~10,000 xg for 15min, followed by 100,000 xg for 30min. The resulting cell lysate was used for affinity purification and flowed over a TALON cobalt affinity resin (Takara Bio). The cobalt resin was washed with ~20 column volumes (CV) of buffer A + 0.1% DDM + 10 mM imidazole, and the protein was subsequently eluted with 4CV of buffer A + 0.1% DDM + 250 mM imidazole. The MsbA sample was then concentrated on a 100 kDa cutoff concentrator and subjected to size-exclusion chromatography on a Superdex 200 column in a 1:1 buffer A water dilution + 0.05% DDM. Peak fractions were pooled and concentrated to ~10 mg/mL for nanodisc reconstitution. MsbA was incubated with membrane scaffold protein MSP1D1 and palmitoyl-oleoyl-phosphatidylglycerol (POPG) lipids at 1:1:60 stoichiometry in size-exclusion buffer + 20 mM sodium cholate. Detergent was removed using ~65% (w/v) biobeads and the sample was again subjected to size-exclusion in 25 mM Tris, 150 mM NaCl, 0.5 mM TCEP. The MsbA nanodisc samples were then concentrated and used for ATPase and cryo-EM experiments.

### ATPase activity assay

MsbA activity was assessed as previously described^1,20,34^. Briefly, TBT1 (DC chemicals) and G247 (Genentech, Inc.) were solubilized and DMSO and used at final DMSO concentration <2%. 1 μg MsbA was pre-incubated in 4°C nanodisc buffer with varying (TBT1, G247, AMP-PNP) inhibitor concentration, then mixed with 2 mM ATP + 2 mM MgCl2. The ATPase reaction was stopped after ~20min at 37°C using 12% sodium dodecyl-sulfate (SDS). γ-phosphate dissociation was assessed in a colorimetric assay by addition of a 6% ascorbic acid (w/v), 1% (w/v) ammonium molybdate in 1N HCl, succeeded by mixing with 2% sodium citrate (w/v), 2% (w/v) sodium meta-arsenite, 2% acetic acid (v/v) in 1N HCl. Absorbance at 850 nm was determined using a Flexstation III Molecular Devices spectrophotometer and data was analyzed on GraphPad Prism 8.

### Negative-stain EM

Protein samples were assessed for homogeneity by negative-stain EM. All samples were stained using a 1.5% (w/v) uranyl formate solution and imaged on a Tecnai T12 electron microscope (FEI), operated at 120kV and at 67,000x magnification. For quantification of MsbA conformations, only clearly inward-facing classes with 2 visible and separate NBDs were considered. The 1mM AMP-PNP/Mg^2+^ sample contained 1 inward-facing class in 30 classes (405 particles in 10,688) and the 1mM AMP-PNP/Mg^2+^ + 5μM G247 contained 18 inward-facing classes in 30 classes (10,784 particles in 17,427).

### Cryo-EM sample preparation and data collection

Both *A. baumannii* and *E. coli* MsbA were first concentrated to ~1.5 mg/mL. Inhibitor-bound complexes were pre-incubated with either 400 μM TBT1 or 100 μM G247 for 2 hours at 4°C, whereas apo *A. baumannii* was directly used for sample preparation. 3 μl nanodisc sample was applied to glow-discharged Quantifoil (R1.2/1.3, 400 mesh) holey carbon grids and blotted with either a Gatan Cryoplunge 3 (blot ~3s) or Thermo Fisher Vitrobot Mark IV (blot ~7s, force +12) before vitrification in liquid ethane. Apo *A. baumannii* and TBT1 -bound *A. baumannii* MsbA were collected on a Titan Krios equipped with a Gatan K3 camera and a BioQuantum imaging filter. Movie stacks were acquired in super-resolution mode with 0.53 Å pixel size and total exposure of 51.6 e/Å^2^. A slit width of 20 eV for energy filter was set during the data collection. The G247-bound *E. Coli* MsbA sample combines data from two microscopes: #1 Titan Krios equipped with a K2 camera (0.42 Å super-resolution pixel size, 62.7 e/Å^2^ exposure) and a BioQuantum imaging filter (with a slit width set to 20 eV); #2 FEI T30 Polara with a K2 camera (0.62 Å super-resolution pixel size, 52 e/Å^2^ exposure). SerialEM^35^ was used for collection of all datasets. More information about cryo-EM data collection parameters can be found in Extended Data Table 1.

### Cryo-EM data processing

The data was analyzed similarly to previous datasets^1,20,34^. Briefly, movie stacks were binned 2x and motion-corrected with MotionCor2^36^, and contrast transfer function was computed with CTFFIND4^37^. All 2D processing was conducted using Simplified Application Managing Utilities of EM Labs (SAMUEL)^1^, which relies on the SPIDER^38^ image processing system. Briefly, ~2,000 protein images were manually picked to generate initial 2D templates. Auto-picking on 10% of movie stacks were then used to create refined 2D templates, then used to pick particles on the whole dataset. 2D particles were screened using “samtree2dv3.py”, which successively runs PCA, *K*-means clustering, and multireference alignment to classify particles. Selected particles after 2D screening were used for 3D processing with RELION3.0^39^, including 3D classification, domain masking, signal subtraction^40^, and 3D particle refinement. Overall resolution of cryo-EM map was computed according to the gold standard FSC method, with reconstruction of independent halves of the dataset and determination of resolution using FSC=0.143 criterion, whereas local resolution was estimated with ResMap^41^. All maps were subjected to density modification^42^ in phenix, with local sharpening and blurring. No model was provided to the density modification program. More information about cryo-EM processing can be found in processing flowcharts (Suppl. Fig. 2,5C,6A).

### Model building and validation

A homology model of *A. baumannii* MsbA was first created with SWISSMODEL^43^ from LPS-bound MsbA (pdb: 5TV4) as input, then manually fit into cryo-EM densities using COOT^44^. G247-MsbA was fit into the cryo-EM map using 5TV4.pdb as reference, then adjusted with COOT. TBT1 and G247 models and restraints were manually built using ELBOW^45^ then fit into the density. The models were then refined against the cryo-EM map using Phenix^46^ real space refinement and validated with MolProbity^47^. All figures were generated with UCSF Chimera^48^ or ChimeraX^49^.

### Structure-based virtual screening and molecular dynamics

A two-step virtual screen was performed to identify compounds binding to the TBT1 pocket. First, flexible docking calculations were carried out for the ChemBridge library (~800,000 compounds) with AutoDock Vina^50^. The center of the TBT1 pocket was selected as coordinate origin, and the docking box set to a 25Å size. Other parameters of Vina were set as default. The next step involved re-scoring of the top 1% docked compounds by an in-house protein-ligand interaction similarity protocol. The BioLip libary^51^, which curates biologically relevant ligand-protein binding interactions, was used to search for similar template pockets as the TBT1 site with PPS-align (availability: https://zhanglab.ccmb.med.umich.edu/PPS-align/). A weighted Jaccard score combining the pocket similarity and ligand similarity was used to rank docked compounds. The top 40 compounds were selected for further validation, and 17 compounds were ordered for experimental testing according to availability. Once W13 was identified as a hit, its binding pose was verified by molecular dynamics (MD) simulation at 310 K and 1 atm. MsbA were embedded into a lipid bilayer consisted of 215 POPC molecules and then solvated into a water box consisted of 31,142 TIP3P water and 0.15 M NaCl. The simulation system included 140,326 atoms and had a dimension of 95 ×95 ×165 Å. The force field parameters of W13 ligand were generated by CGenFF^52^, and the protein was modeled by the CHARMM36m force field^53^. System setup and analysis were performed with CHARMM^54^ and MD simulations were carried out using OpenMM^55^.

---

**Supplementary Figure 1.**
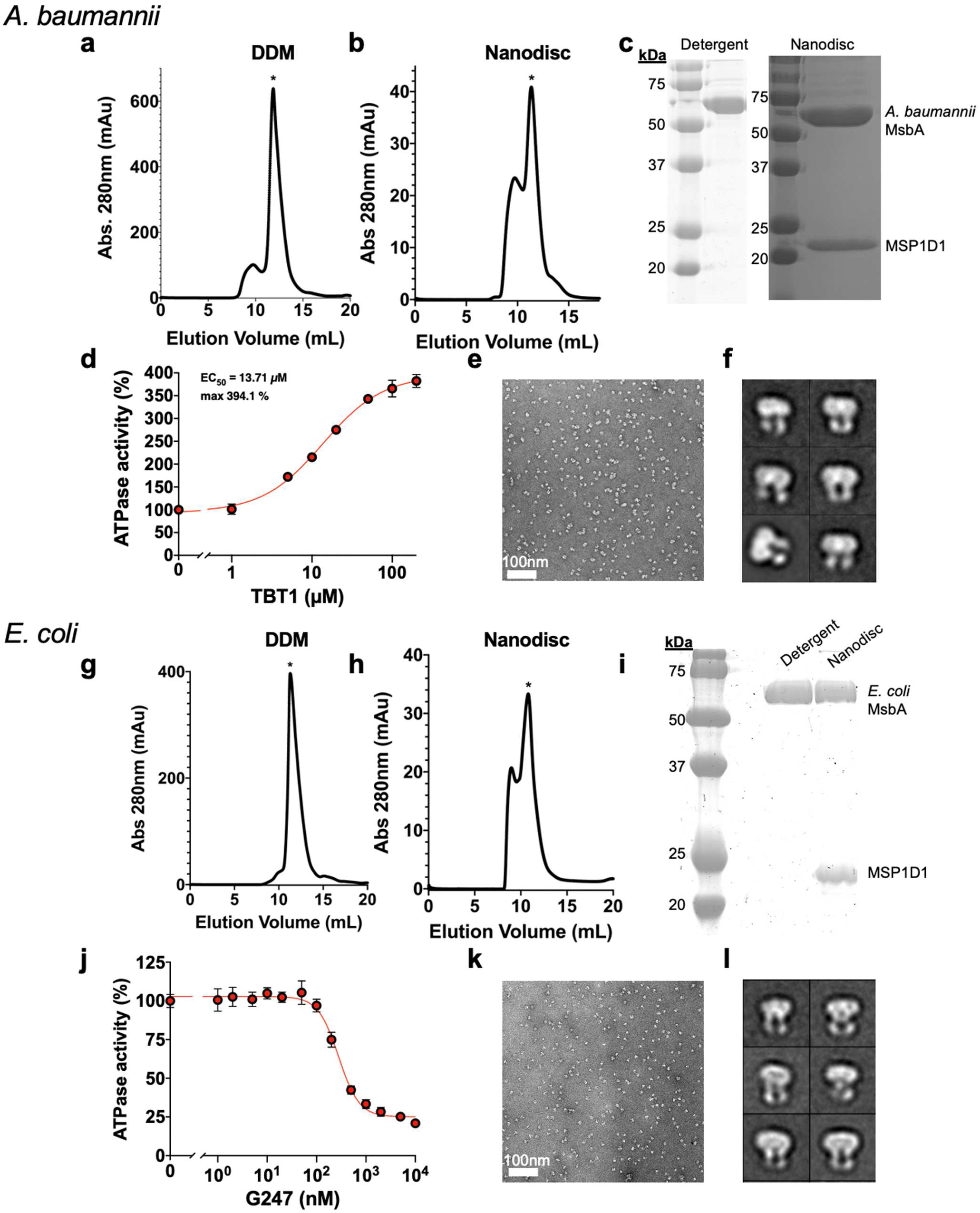
Characterization of *A. baumannii* and *E. coli* MsbA. **a,** Gel filtration profile of purified *A. baumannii* MsbA in DDM, ran on a Superdex 200 column. The protein elution peak is marked with a star symbol. **b,** Gel filtration profile of *A. baumannii* MsbA in MSP1D1 nanodiscs. Incorporated MsbA is marked with a star symbol. **c,** SDS-PAGE gels of *A. baumannii* MsbA in DDM (left) and incorporated in nanodiscs (right). **d,** ATPase activity of *A. baumannii* MsbA when exposed to increasing TBT1 concentrations. Each point represents mean ± SD (n=3). **e,** Representative negative-stain EM image of *A. baumannii* MsbA in nanodiscs. **f,** Representative negative-stain 2D class averages of *A. baumannii* MsbA in nanodiscs. Box size is 215 Å. **g,** Gel filtration profile of *E. coli* MsbA in DDM detergent. **h,** Gel filtration profile of *E. coli* MsbA in nanodiscs. **i,** SDS-PAGE analysis of purified *E. coli* MsbA after gel filtration. **j,** ATPase activity of *E. coli* MsbA with increasing G247 concentration. Each point represents mean ± SD (n=3). **k,** Negative-stain EM image of *E. coli* MsbA. **l,** Negative-stain 2D class averages of *E. coli* MsbA in nanodiscs, with a 215 Å box size.

**Supplementary Figure 2.**
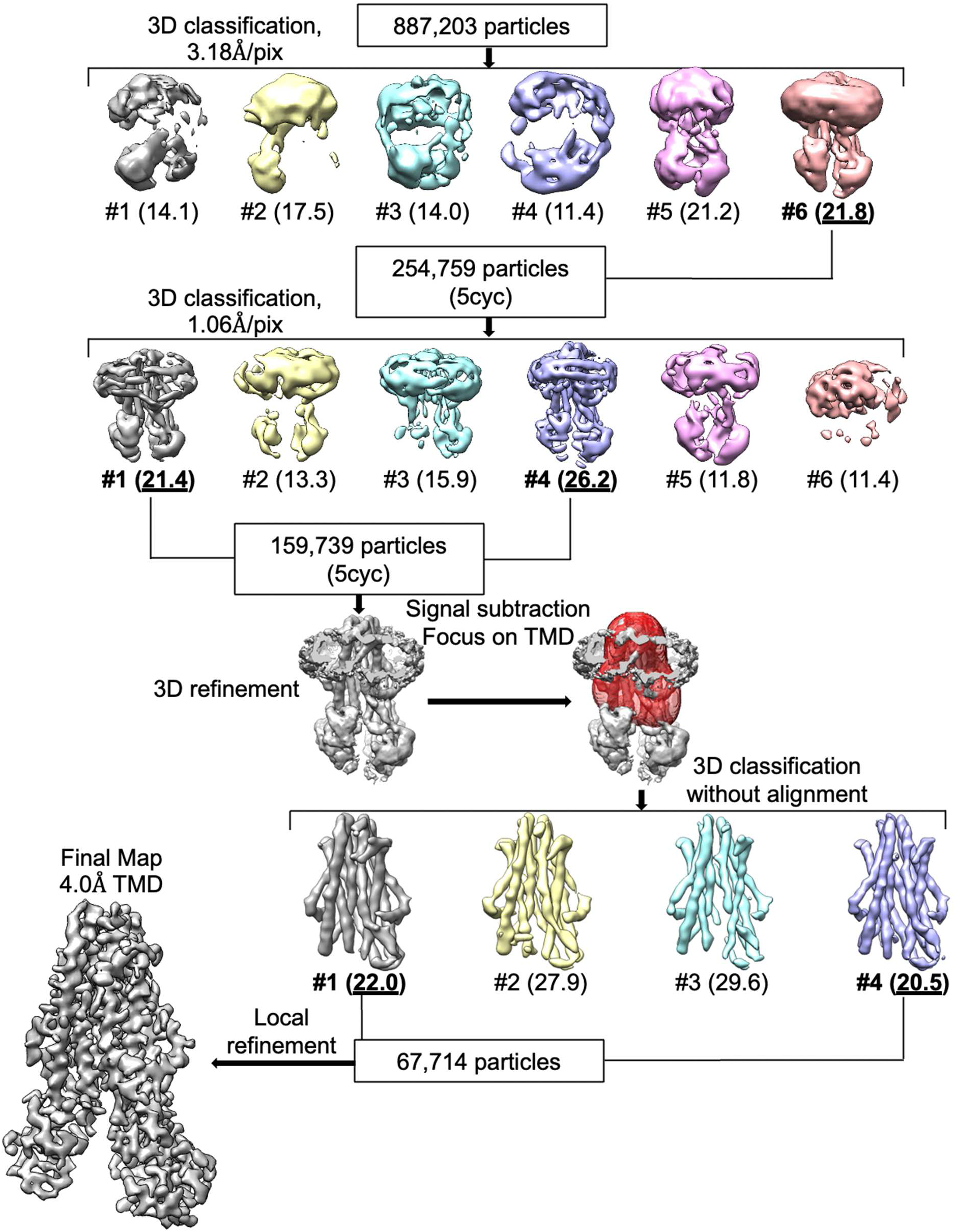
Processing flowchart for *A. baumannii* MsbA in complex with TBT1. “5cyc” refers to the method of including any particle being assigned to a selected class within the final 5 cycles in 3D classification.

**Supplementary Figure 3.**
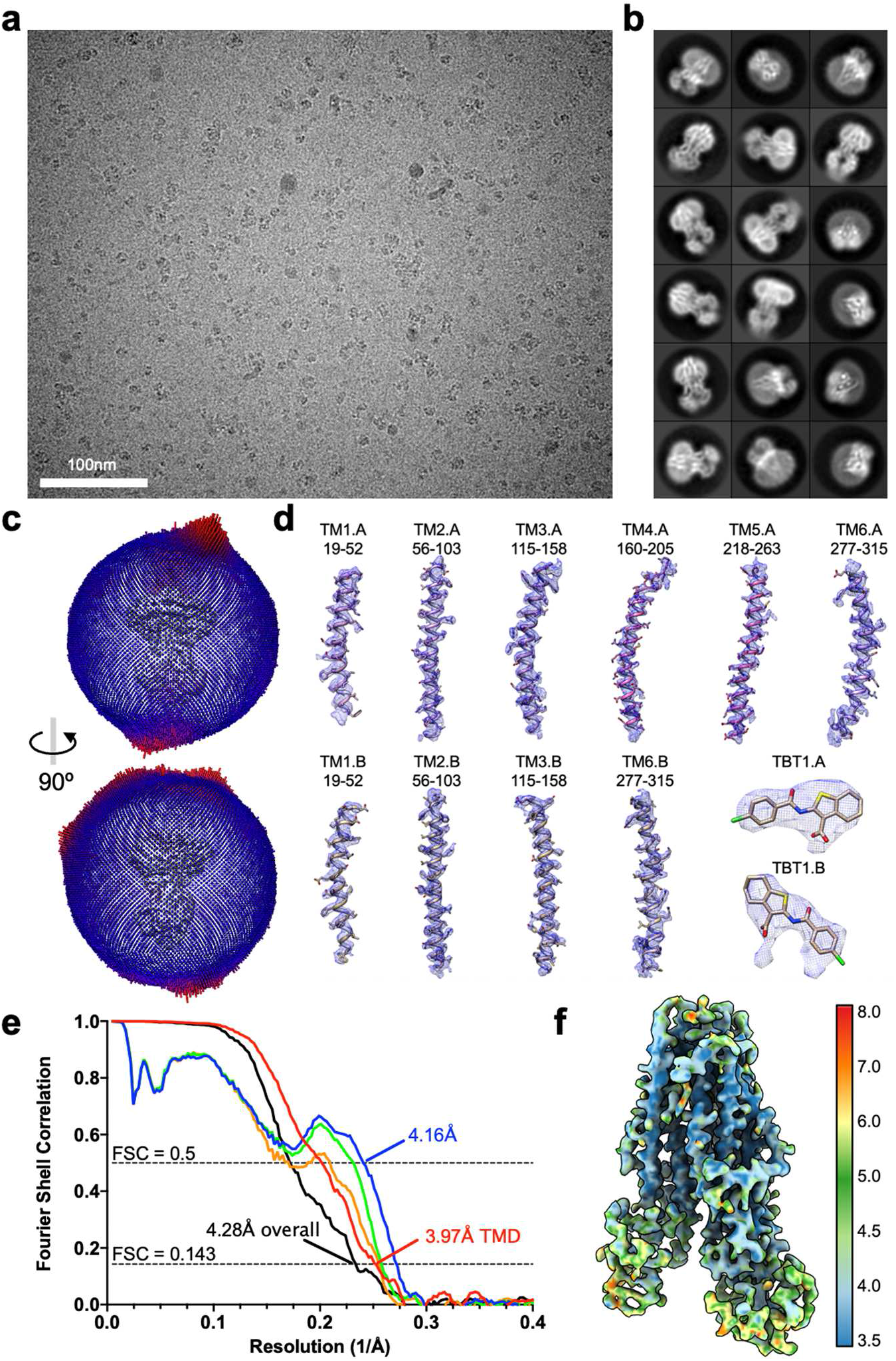
Cryo-EM imaging and validation for A. *baumannii* MsbA in complex with TBT1. **a,** Representative cryo-EM image of TBT1-bound MsbA. **b,** 2D class averages of *A. baumannii* MsbA bound to TBT1. The asymmetric positioning of NBDs is readily discernable. Box size is 203 Å. **c,** Angle distribution of the cryo-EM particles included for final 3D reconstruction. **d,** Close-up view of all ordered transmembrane helices and TBT1 densities. **e,** Fourier Shell Correlation (FSC) curves of the final reconstruction. Half map #1 vs. half map #2 for the entire MsbA molecule is shown in black. The remaining FSC curves were calculated for TMDs only: half map #1 vs. half map #2 (red), model vs. refined map (blue), model refined in half map #1 vs. half map #1 (green), and model refined in half map #1 vs. half map #2 (orange). **f,** Local resolution of the final cryo-EM map.

**Supplementary Figure 4.**
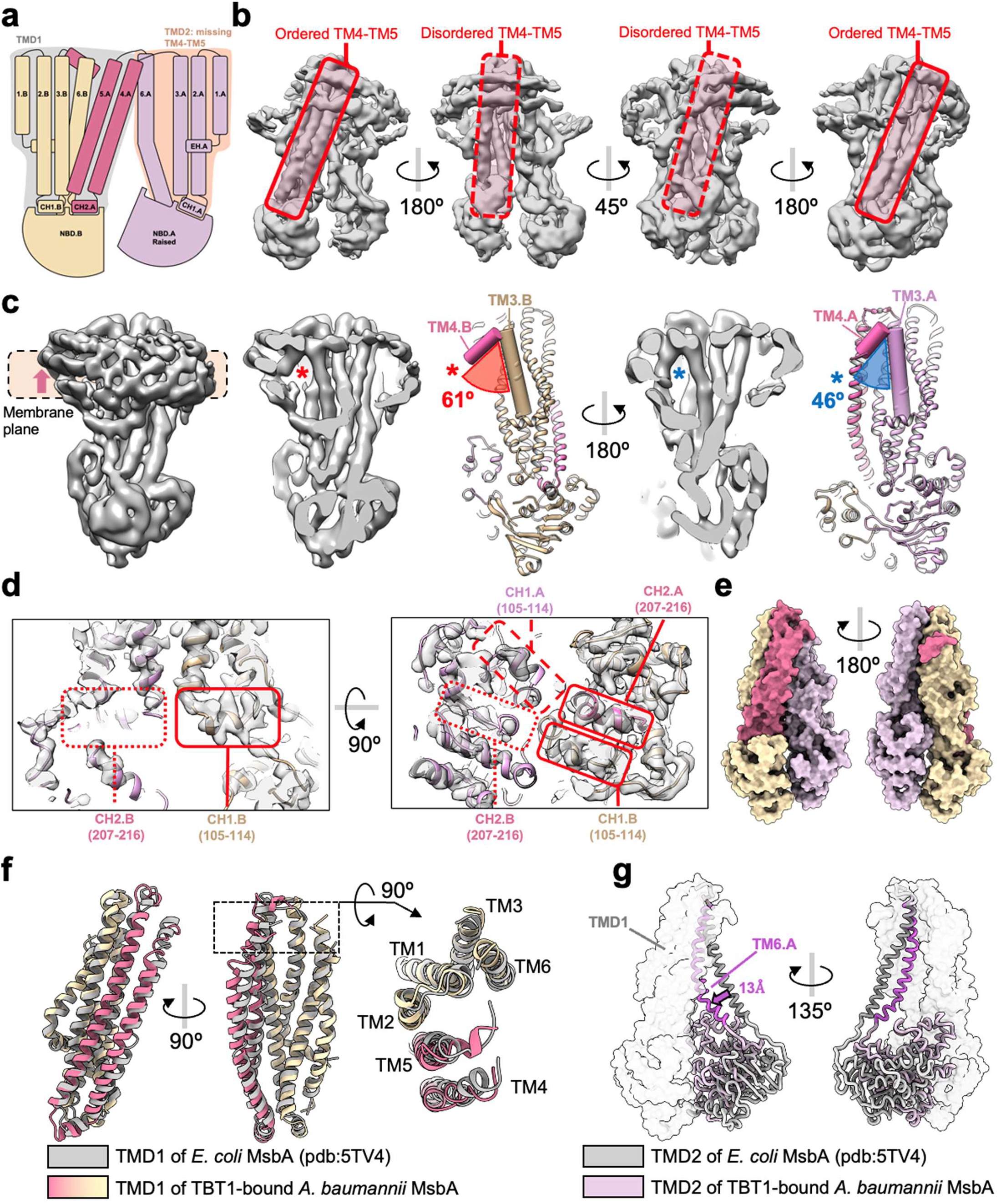
Structural analysis of TBT1-inhibited MsbA. **a,** Reproduction of TBT1-bound MsbA cartoon from Fig. 1c. **b,** Rotated side views of the unsharpened, unmasked cryo-EM map of TBT1-bound MsbA. The location of the domain-swapping TM4-TM5 bundles is highlighted in red box, with solid line when ordered and dashed line when disordered. **c,** Angle comparison between TM3 and the N-terminal end of TM4 for each MsbA chain. In chain B (left, red asterisk), TM4 is kinked outward and pushes the nanodisc scaffold upwards and away from the membrane plane. In chain A (right, blue asterisk), the separation between TM3 and TM4 is narrower. For angle measurements, axes were defined using the “define axis” function in UCSF chimera and selecting residues 133-157 for TM3 and 158-165 for TM4. **d,** Map and model highlighting the location of the coupling helices (CH). Coupling helices are boxed with a solid line when present, dashed line when destabilized, and dotted line when disordered. **e,** Surface representation of the model of TBT1-bound MsbA, highlighting the constricted TMDs. **f,** Alignment of the TMD1 bundle of TBT1-bound *A. baumannii* MsbA (colored) with drug-free *E. coli* MsbA (PDB: 5TV4, gray). **g,** Different positioning of TM6.A between TBT1-bound *A. baumannii* MsbA (pink) and drug-free *E. coli* MsbA (PDB: 5TV4, gray). The structurally similar TMD1 is shown as transparent surface, while TM6.A and NBD.A are displayed using worm representation. TM6.A is colored in darker pink or gray to highlight its location. The 13 Å shift of TM6 is measured using Ala308 as reference.

**Supplementary Figure 5.**
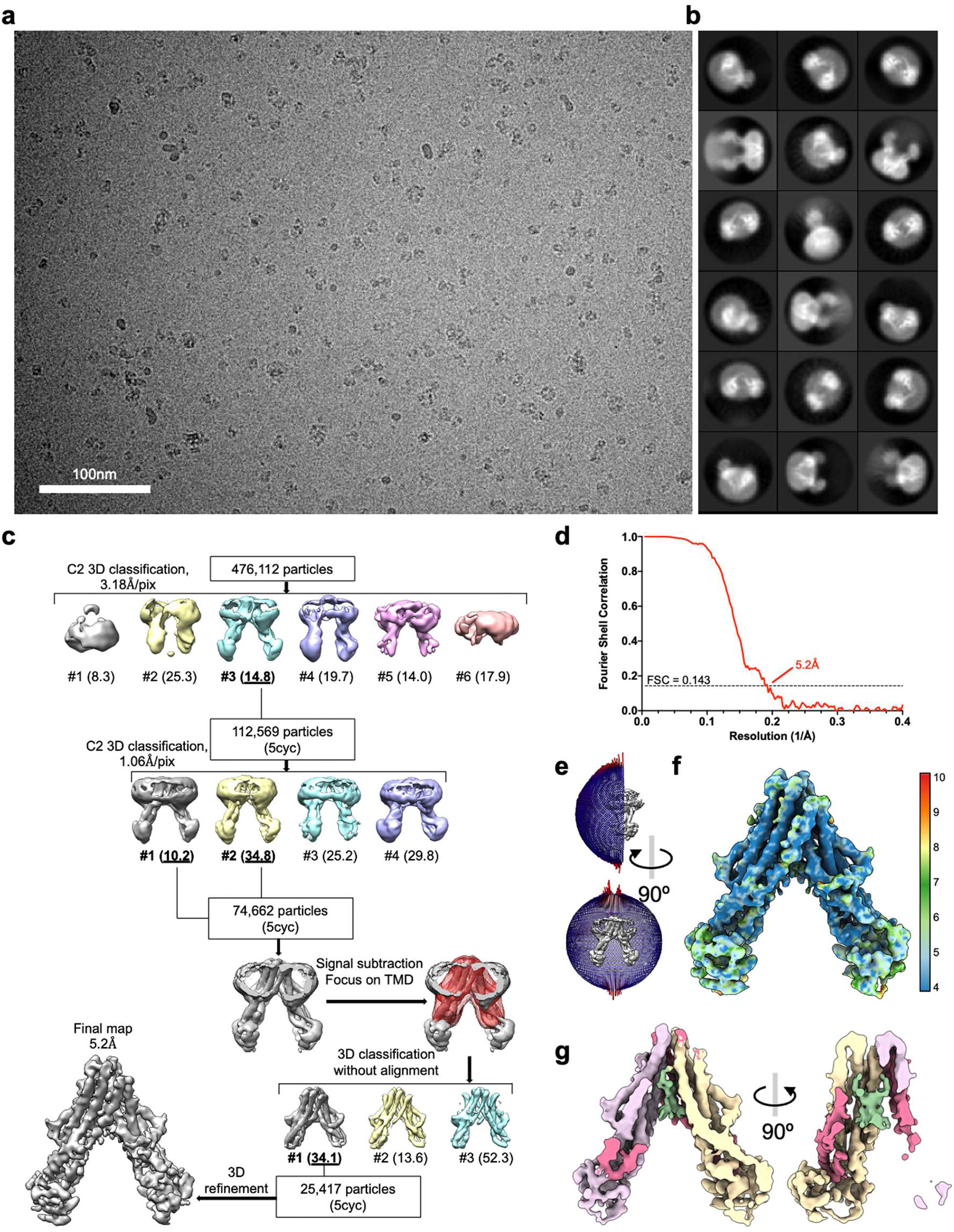
Cryo-EM imaging and validation for drug-free A. *baumannii* MsbA. **a,** Representative cryo-EM image of MsbA in nanodiscs in the absence of compound. **b,** 2D class averages of *A. baumannii* MsbA. Box size is 203 Å. **c,** Processing flowchart for the cryo-EM dataset. **d,** FSC curve of independently refined half-maps for the final refinement. **e,** Angle distribution of the cryo-EM particles included for final 3D reconstruction. **f,** Local resolution for the final cryo-EM map. **g,** A C2-symmetrized LPS-like density, colored dark green, is present in the central substrate-binding pocket.

**Supplementary Figure 6.**
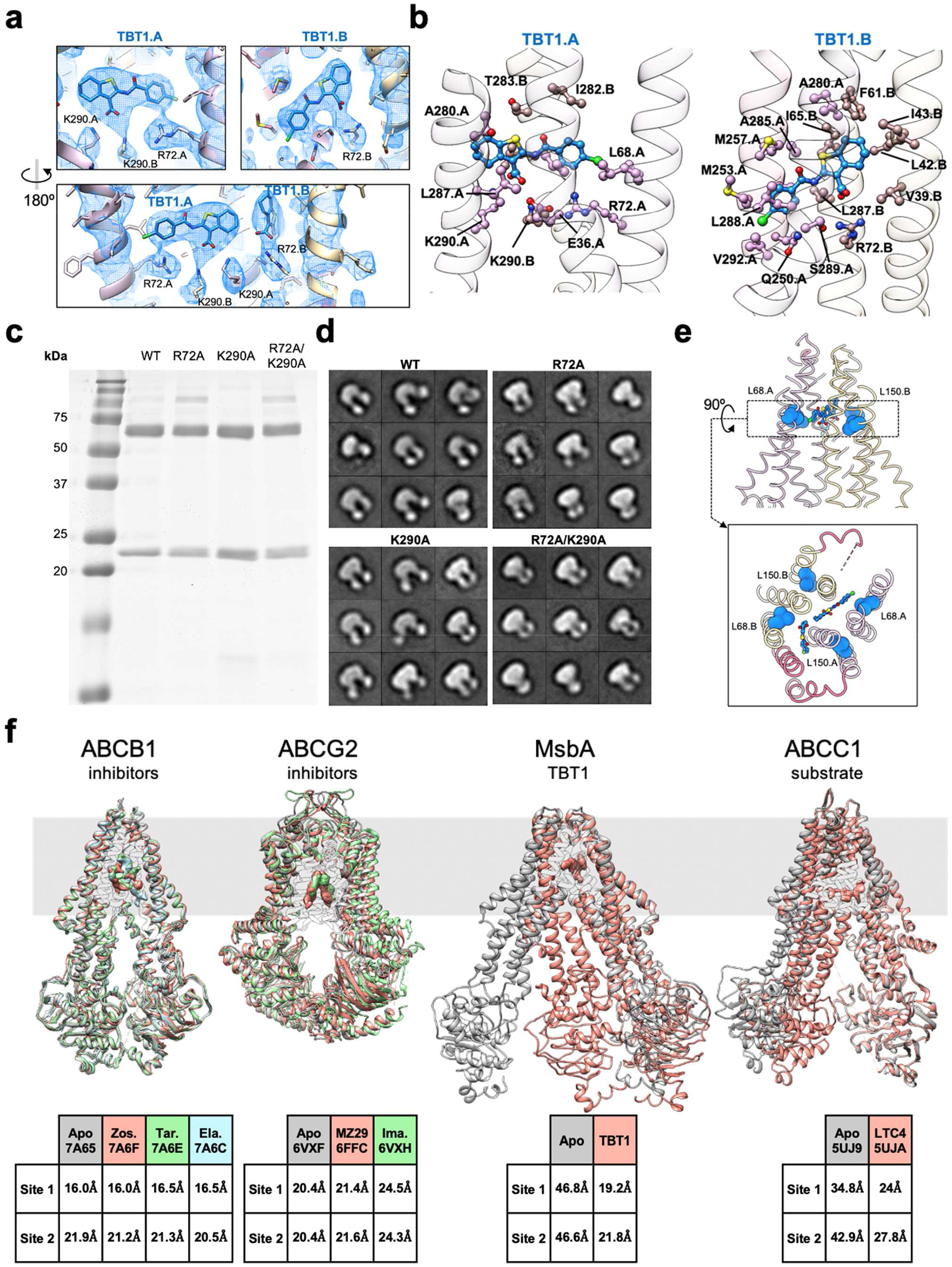
Molecular details of TBT1 binding to *A. baumannii* MsbA. **a,** Superimposition of cryo-EM map and model of the TBT1 binding site. **b,** Binding pocket for each asymmetric TBT1 ligand. **c,** SDS-PAGE gel of purified MsbA proteins in nanodiscs. *A. baumannii* MsbA runs at ~65 kDa and MSP1D1 at ~22.5 kDa. **d,** Negative-stain 2D class averages of wild-type and mutant *A. baumannii* MsbA. Box size is 215 Å. **e,** Location of mutations (blue spheres) previously found to prevent TBT1-induced ATPase stimulation and LPS transport blockage. **f,** Comparison of various cryo-EM structures of apo and compound-bound ABC transporters. Top: alignment of apo structures with inhibited and substrate-bound conformations. Ligands are shown as zoned densities from deposited cryo-EM maps. The multidrug efflux pump ABCB1 adopts a similar conformation whether in apo state (PDB: 7A65) or inhibited by zosuquidar (“zos.”, PDB: 7A6F), tariquidar (“tar.”, PDB: 7A6E) and elacridar (“ela.”, PDB: 7A6C). ABCG2 (PDB: 6VXF) undergoes modest conformational changes in complex with its inhibitors MZ29 (PDB: 6FFC) and imatinib (“ima.”, PDB: 6VXH). In contrast, TBT1-bound MsbA undergoes a large-scale conformational change, which is reminiscent of NBD tightening when ABCC1 (PDB: 5UJ9) is exposed to its LTC4 endogenous substrate (PDB: 5UJA). Bottom: Distance between Cα signature motif serine and Walker A glycine for each ATP binding site. For ABCB1, site 1 is S532/G1075 and site 2 is G432/S1177. For ABCG2, S187/G85 is used. For MsbA, S474/G373. For ABCC1, site 1 is S769/G1329 and site 2 is S1430/G681.

**Supplementary Figure 7.**
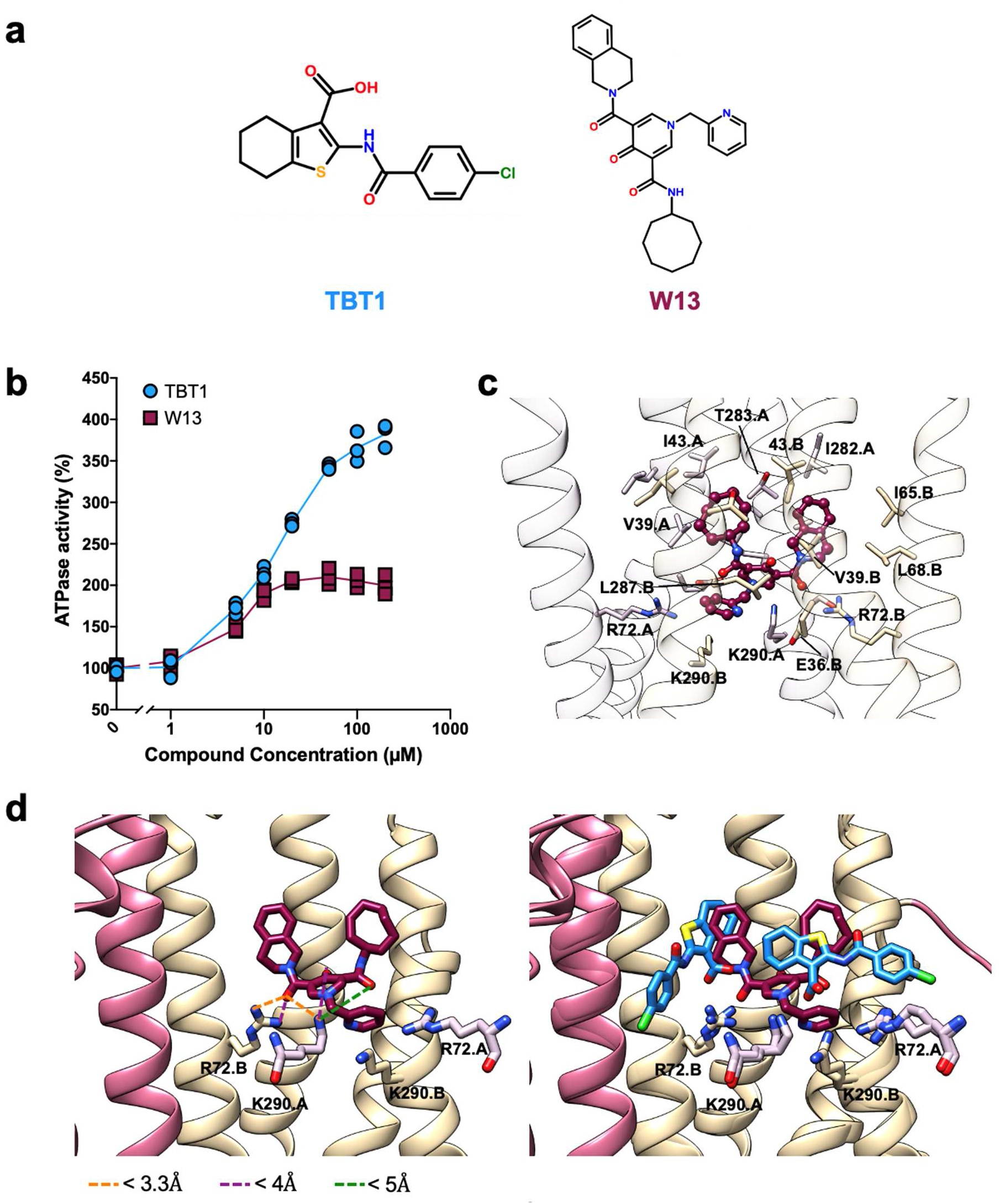
Predicted binding of W13 to A. *baumannii* MsbA. **a,** Molecular structures of TBT1 and W13. **b,** Effect of TBT1 and W13 on MsbA ATPase activity at different compound concentrations. **c,** W13 binding pocket in the central MsbA cavity, as predicted after 40-ns molecular dynamics. W13 is shown maroon using ball-and-stick representation. Neighboring residues are displayed as sticks with coloring matching their chain assignment. **d,** Predicted W13 fit, with neighboring lysine and arginine residues and distances between sidechains and compound (left). Superimposition of TBT1 (blue) and W13 (maroon) binding pockets (right).

**Supplementary Figure 8.**
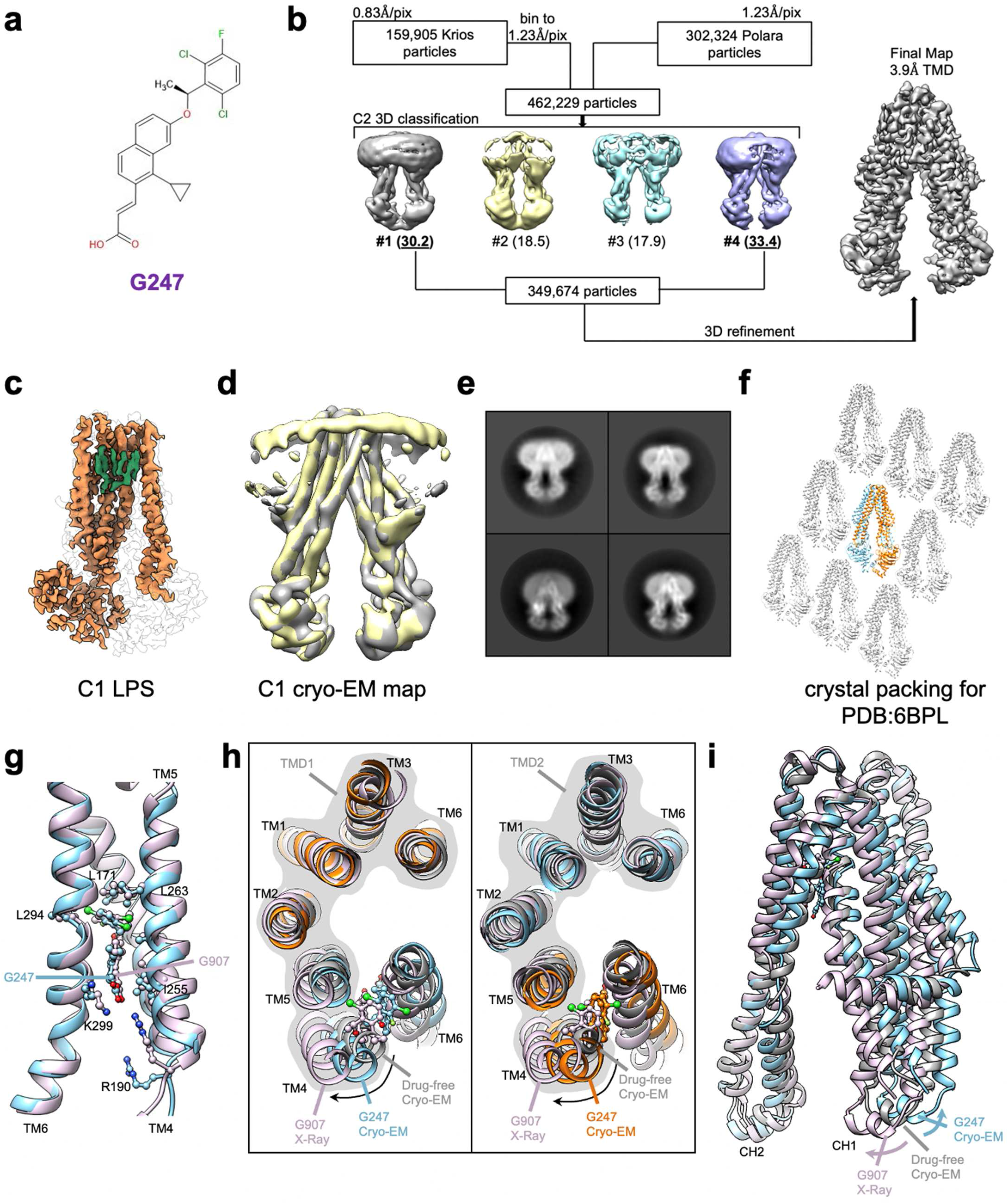
Processing flowchart and analysis for *E. coli* MsbA bound to G247. **a,** Molecular structure of G247. **b,** Processing flowchart for MsbA – G247 dataset. 3D classification and final refinement were performed with C2 symmetry as no apparent asymmetric classes were identifiable from the dataset. **c,** LPS density (green) is clearly resolved in the inner pocket. **d,** The C1-refined map (gray) is superimposed with the same map after 180° rotation (yellow), showing nearly perfect symmetrical structure. **e,**2D class averages of the cryo-EM particle images of G247 bound MsbA, exhibiting symmetrical NBDs. Box size is 236 Å. **f,** Crystal packing of MsbA in complex with G907 (PDB:6BPL). **g,** Comparison of drug-binding pocket for G247 (blue) and G907 (pink, PDB:6BPL), as seen from the LPS binding site. Alignment performed on TM4,5,6. **h,** Helix positioning for drug-free MsbA (gray, PDB:5TV4), G247-bound MsbA (blue and orange), and G907-bound crystal structure (pink, PDB:6BPL). Alignment performed on each TMD, shown with gray background. **i,** Comparison of coupling helices (CH) positioning for drug-free MsbA (gray, PDB:5TV4), G247-bound MsbA (blue), and G907-bound crystal structure (pink, PDB:6BPL). Compared to the reference drug-free cryo-EM structure, the distance between CH1 and CH2 is reduced for the G907-bound crystal structure and increased for G247-bound MsbA. Only one TM chain is shown for clarity. Alignment is performed on TMD1, as shown in panel h.

**Supplementary Figure 9.**
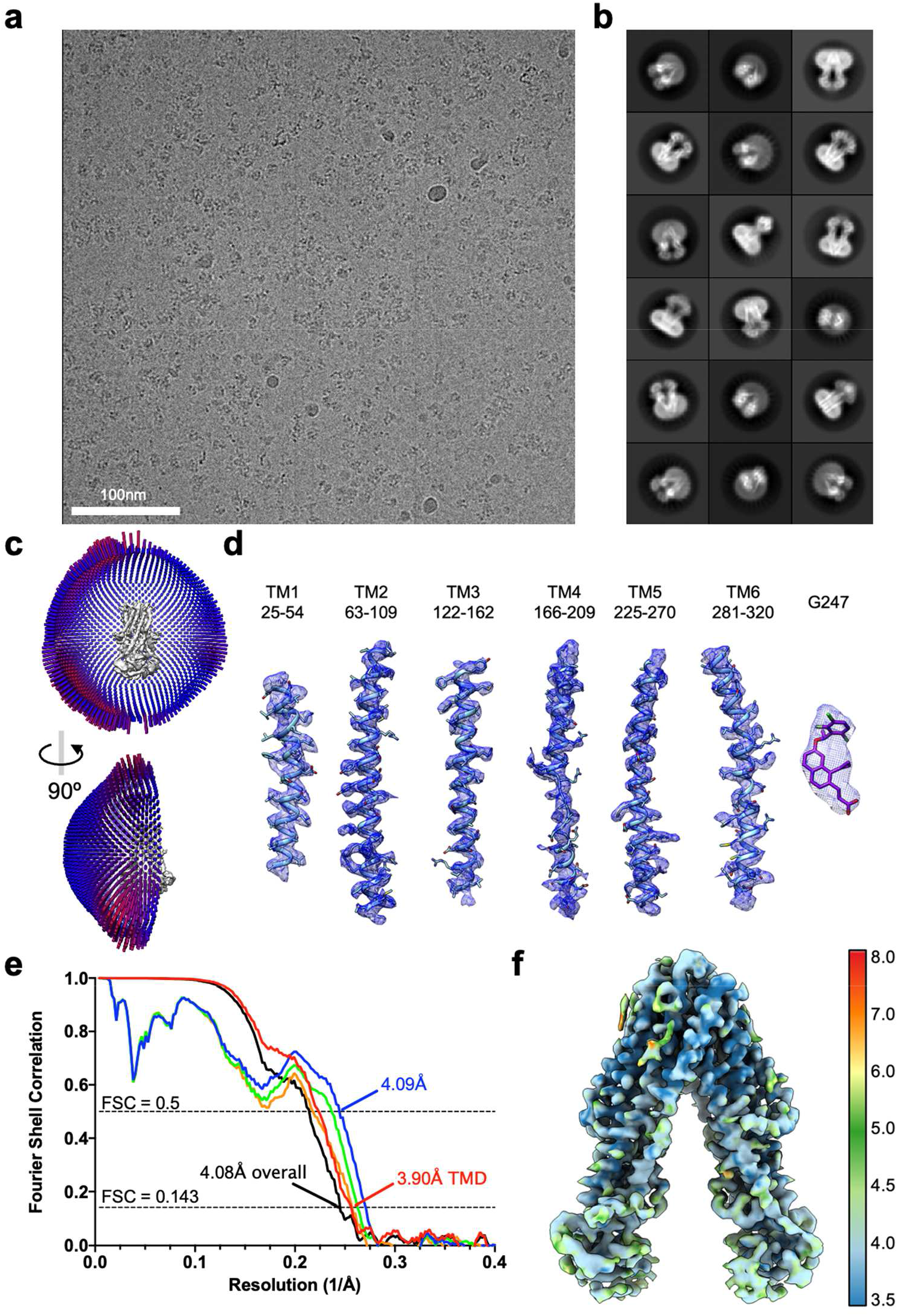
Cryo-EM imaging and validation for *E. coli* MsbA bound to G247. **a,** Representative cryo-EM image of *E. coli* MsbA in complex with G247. **b,** 2D class averages. Box size is 236 Å. **c,** Angle distribution of cryo-EM particles included for the final 3D reconstruction. **d,** Superimposition of cryo-EM density and model of TM helices and G247. **e,** Fourier Shell Correlation (FSC) curves of the final reconstruction. Half map #1 vs. half map #2 for the entire MsbA molecule is shown in black. The remaining FSC curves were calculated for TMDs only: half map #1 vs. half map #2 (red), model vs. refined map (blue), model refined in half map #1 vs. half map #1 (green), and model refined in half map #1 vs. half map #2 (orange). **f,** Local resolution of the final cryo-EM map.

**Supplementary Table 1.**
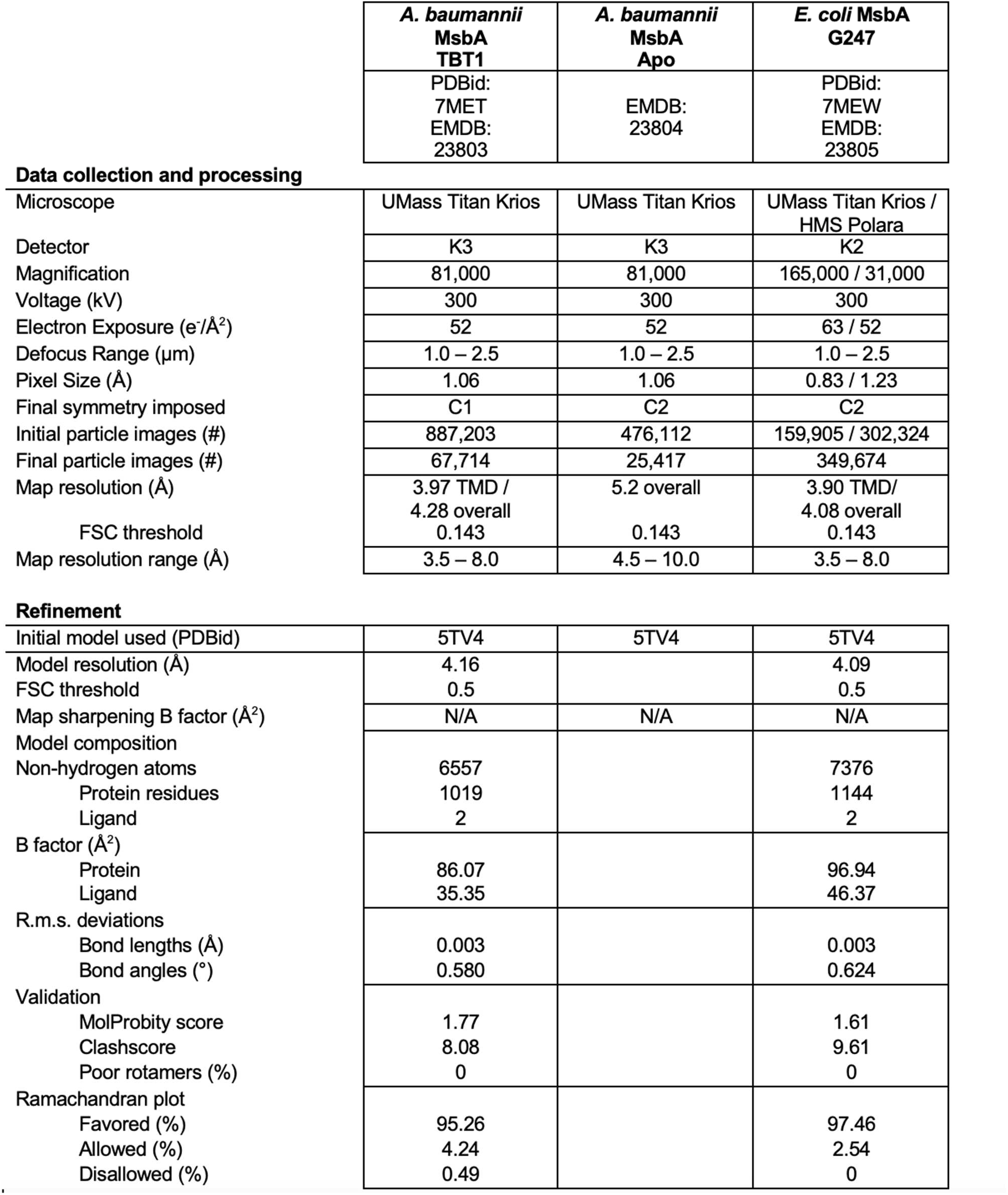
Data collection table and model statistics.

| *A. baumannii*<br>MsbA<br>TBT1 | *A. baumannii*<br>MsbA<br>Apo | *E. coli* MsbA<br>G247 |
| --- | --- | --- |
| PDBid:<br>7MET<br>EMDB:<br>23803 | EMDB:<br>23804 | PDBid:<br>7MEW<br>EMDB:<br>23805 |

Data collection and processing
|  |  |  |  |
| --- | --- | --- | --- |
| Microscope | UMass Titan Krios | UMass Titan Krios | UMass Titan Krios /<br>HMS Polara |
| Detector | K3 | K3 | K2 |
| Magnification | 81,000 | 81,000 | 165,000 / 31,000 |
| Voltage (kV) | 300 | 300 | 300 |
| Electron Exposure (e<sup>-</sup>/Å<sup>2</sup>) | 52 | 52 | 63 / 52 |
| Defocus Range (μm) | 1.0 – 2.5 | 1.0 – 2.5 | 1.0 – 2.5 |
| Pixel Size (Å) | 1.06 | 1.06 | 0.83 / 1.23 |
| Final symmetry imposed | C1 | C2 | C2 |
| Initial particle images (#) | 887,203 | 476,112 | 159,905 / 302,324 |
| Final particle images (#) | 67,714 | 25,417 | 349,674 |
| Map resolution (Å) | 3.97 TMD /<br>4.28 overall | 5.2 overall | 3.90 TMD/<br>4.08 overall |
| FSC threshold | 0.143 | 0.143 | 0.143 |
| Map resolution range (Å) | 3.5 – 8.0 | 4.5 – 10.0 | 3.5 – 8.0 |

**Supplementary Table 1 | Data collection table and model statistics**
|  |  |  |  |
| --- | --- | --- | --- |
| Initial model used (PDBid) | 5TV4 | 5TV4 | 5TV4 |
| Model resolution (Å) | 4.16 |  | 4.09 |
| FSC threshold | 0.5 |  | 0.5 |
| Map sharpening B factor (Å<sup>2</sup>) | N/A | N/A | N/A |
| Model composition |  |  |  |
| Non-hydrogen atoms | 6557 |  | 7376 |
| Protein residues | 1019 |  | 1144 |
| Ligand | 2 |  | 2 |
| B factor (Å<sup>2</sup>) |  |  |  |
| Protein | 86.07 |  | 96.94 |
| Ligand | 35.35 |  | 46.37 |
| R.m.s. deviations |  |  |  |
| Bond lengths (Å) | 0.003 |  | 0.003 |
| Bond angles (°) | 0.580 |  | 0.624 |
| Validation |  |  |  |
| MolProbity score | 1.77 |  | 1.61 |
| Clashscore | 8.08 |  | 9.61 |
| Poor rotamers (%) | 0 |  | 0 |
| Ramachandran plot |  |  |  |
| Favored (%) | 95.26 |  | 97.46 |
| Allowed (%) | 4.24 |  | 2.54 |
| Disallowed (%) | 0.49 |  | 0 |

**Supplementary Table 2.**
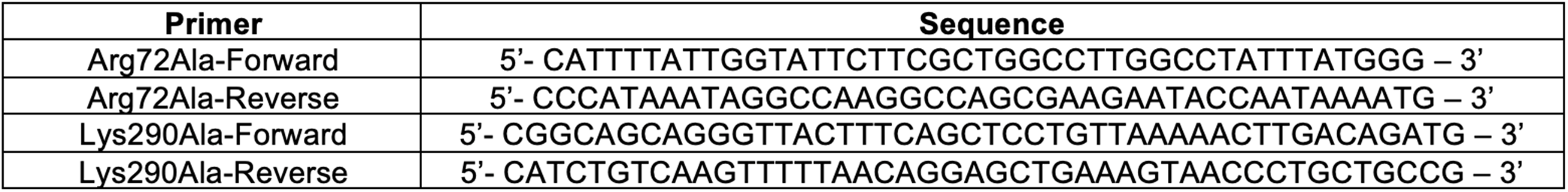
Mutagenesis primers used in this study.

